# Discovery of a cell-active chikungunya virus nsP2 protease inhibitor using a covalent fragment-based screening approach

**DOI:** 10.1101/2024.03.22.586341

**Authors:** Eric M. Merten, John D. Sears, Tina M. Leisner, Paul B. Hardy, Anirban Ghoshal, Mohammad Anwar Hossain, Kesatebrhan Haile Asressu, Peter J. Brown, Michael A. Stashko, Laura Herring, Angie L. Mordant, Thomas S. Webb, Christine A. Mills, Natalie K. Barker, Zachary J. Streblow, Sumera Perveen, Cheryl Arrowsmith, Jamie J. Arnold, Craig E Cameron, Daniel N. Streblow, Nathaniel J. Moorman, Mark Heise, Timothy M. Willson, Konstantin Popov, Kenneth H. Pearce

## Abstract

Chikungunya virus (CHIKV) is a mosquito-borne alphavirus that has been responsible for numerous large-scale outbreaks in the last twenty years. Currently, there are no FDA-approved therapeutics for any alphavirus infection. CHIKV non-structural protein 2 (nsP2), which contains a cysteine protease domain, is essential for viral replication, making it an attractive target for a drug discovery campaign. Here, we optimized a CHIKV nsP2 protease (nsP2pro) biochemical assay for the screening of a 6,120-compound cysteine-directed covalent fragment library. Using a 50% inhibition threshold, we identified 153 hits (2.5% hit rate). In dose-response follow up, RA-0002034, a covalent fragment that contains a vinyl sulfone warhead, inhibited CHIKV nsP2pro with an IC_50_ of 58 ± 17 nM, and further analysis with time-dependent inhibition studies yielded a k_inact_/K_I_ of 6.4 x 10^3^ M^-1^s^-1^. LC-MS/MS analysis determined that RA-0002034 covalently modified the catalytic cysteine in a site-specific manner. Additionally, RA-0002034 showed no significant off-target reactivity against a panel of cysteine proteases. In addition to the potent biochemical inhibition of CHIKV nsP2pro activity and exceptional selectivity, RA-0002034 was tested in cellular models of alphavirus infection and effectively inhibited viral replication of both CHIKV and related alphaviruses. This study highlights the discovery and characterization of the chemical probe RA-0002034 as a promising hit compound from covalent fragment-based screening for development toward a CHIKV or pan-alphavirus therapeutic.

**Significance Statement:** Chikungunya virus is one of the most prominent and widespread alphaviruses and has caused explosive outbreaks of arthritic disease. Currently, there are no FDA-approved drugs to treat disease caused by chikungunya virus or any other alphavirus-caused infection. Here, we report the discovery of a covalent small molecule inhibitor of chikungunya virus nsP2 protease activity and viral replication of four diverse alphaviruses. This finding highlights the utility of covalent fragment screening for inhibitor discovery and represents a starting point towards the development of alphavirus therapeutics targeting nsP2 protease.

## Introduction

Alphaviruses are widespread, enveloped, single-stranded positive sense RNA viruses transmitted by *Aedes aegypti* and *Aedes albopictus* mosquitos and pose a serious threat to public health(1). Alphaviruses are separated into two classes based on geographical emergence: Old World alphaviruses such as chikungunya virus (CHIKV), Ross River virus (RRV), and O’nyong O’nyong virus (ONNV), and New World alphaviruses, which include Venezuelan (VEEV), Western (WEEV) and Eastern (EEEV) Equine Encephalitis viruses and Mayaro virus (MAYV). Old World alphavirus infections are characterized by rash, fever, and in some cases prolonging arthralgia which can persist for months post-infection. Rarer symptoms such as neurological complications, severe fever, and death can occur in vulnerable populations including young children and the elderly (2). The clinical presentations of New World alphaviruses, which are distinct from Old World alphaviruses, more commonly cause neurological symptoms such as encephalitis, along with fever, headache, and nausea(3). New World alphavirus infections can be deadly, specifically EEEV, with 30-50% of EEEV cases resulting in mortality. In 2019, there were record high cases of EEEV in the United States with 34 cases and 12 deaths, increased significantly from the previous median of 8 cases per year(4).

CHIKV, one of the most prevalent and widely distributed alphaviruses, is currently a serious public health concern. CHIKV was first identified in Eastern Africa in 1952 and has since spread globally with cases reported in over 100 countries(5). CHIKV is now endemic to Southeast Asia, Central Africa, Pacific, Caribbean, and Central and South America. CHIKV first emerged in the Americas in 2013 on the island of St. Martin, and by 2016 close to one million cases had been reported(6). Currently, there is an ongoing CHIKV epidemic in Paraguay with ∼120,000 cases reported between October 2022 and April 2023(7). The geographical expansion of CHIKV has been driven by mutational and environmental factors. Historically, CHIKV was transmitted by *Aedes aegypti* mosquitos. However, an adaptive mutation on the E1 envelope protein (A226V) caused increased fitness for *Aedes albopictus* resulting in a notable outbreak on the island of Réunion in 2005 wherein over one third of the island’s population contracted CHIKV (∼250,000 cases)(8). Additionally, the habitat for *Aedes sp*. mosquitos is expanding. Recently, reports of Dengue virus, another arbovirus transmitted by *Aedes sp*. mosquitos, was found at altitudes previously uninhabitable by *Aedes sp*. mosquitos(9). According to computational models, the changing climate has the potential to expose up to one billion people to new viral transmission in the next 50 years(10). As of 2023, there are no current FDA-approved drugs for any alphavirus-caused disease. The recommended treatments for CHIKV are non-steroidal anti-inflammatory drugs as well as mosquito population control, highlighting the unmet medical need for development of CHIKV and alphavirus therapeutics(11).

Direct-acting antivirals, which target the viral proteins essential for viral replication, have yet to be reported for therapeutic use against CHIKV. The CHIKV genome contains two open-reading frames (ORFs) that encode two polyproteins; the first polyprotein (P1234) contains machinery necessary for replication and transcription (nsP1-4) and the second polyprotein contains the structural components to form new virions (Capsid, E3, E2, 6K, E1)(1). Polyprotein P1234 is autoproteolyzed by non-structural protein 2 (nsP2) to form four individually active nsPs, 1-4. nsP2 is the largest non-structural protein and contains an N-terminal RNA-specific helicase domain (nsP2hel) and a C-terminal cysteine protease domain (nsP2pro) and methyl-transferase-like domain. The papain-like nsP2pro domain contains the papain catalytic dyad of cysteine (C478) and histidine (H548) residues, however nsP2pro lacks an essential catalytic aspartate residue found in papain and shares little structural similarity with papain outside of the two catalytic residues. CHIKV nsP2pro is an attractive drug target because nsP2pro proteolytic activity is essential for viral replication(12). A series of compounds developed using computer-aided drug design with weak inhibitory activity against CHIKV nsP2pro have been reported, however these compounds lack experimentally confirmed mechanisms of inhibition and target engagement data necessary for advancement (13–15). VEEV inhibitors have been identified that either inhibit VEEV nsP2pro or are tracked to nsP2 through resistance mutations(16–19). However, none of these inhibitors has been shown to inhibit CHIKV nsP2pro or potently inhibit CHIKV replication. In addition to the lack of drug candidates targeting nsP2pro, there is a notable lack of tool compounds that can be used for the broad study of alphaviruses and alphavirus nsP2pro. These tool compounds or chemical probes are selective small-molecule modulators that can be utilized to interrogate and validate target function in a biological system(20). A high-quality chemical probe for CHIKV nsP2pro can also serve as a starting point for development of an alphavirus therapeutic.

One strategy for small molecule protease inhibition is the covalent modification of amino acids involved in catalysis with targeted covalent inhibitors (TCIs) (21). TCIs are compounds that contain a reactive functional group (a.k.a. ‘warhead’) which require selective compound binding to a location in proximity to a nucleophilic residue (Cys, Lys, Ser, etc.), thus enabling a binding-induced covalent adduct. Cysteines are the most common nucleophilic residues modified by TCIs(22). CHIKV nsP2pro is a strong candidate to target with TCIs due to the essential nature of the active-site C478 for enzymatic function and viral replication. Studies have shown that C478 cannot be replaced with a neighboring residue for catalysis, as C478A mutations abolish protease activity of the protein(23). Historically, TCIs have been avoided in the drug discovery pipeline due to potential liabilities such as toxicity and off-target reactivity with glutathione (GSH) or other macromolecules. In the last decade however, interest in TCIs as therapeutics and chemical probes has increased due to the benefits of irreversible inhibition that include increased biochemical potency and longer target residency times compared to reversible inhibitors(24). Many safe, widely used medications exploit a covalent mechanism of action, including aspirin and penicillin(25, 26). Viral proteases from other virus families have also been successfully targeted with TCIs, with FDA-approved drugs for SARS-CoV-2 M_pro_ (nirmatrelvir) and HCV NS3 (boceprevir, since discontinued), which both covalently target nucleophilic catalytic residues(27, 28).

TCI development often begins with an existing non-covalent inhibitor for which an electrophile is incorporated into the structure to induce covalent bond formation with a nearby nucleophilic residue. However, there are currently no reported, validated, active site-directed inhibitors that bind CHIKV nsP2pro in proximity to C478. An alternative strategy for discovery of TCIs is via covalent fragment-based drug discovery (CFBDD), in which a small fragment (<300 Da) is tethered to an electrophilic warhead and screened against the target of interest (29, 30). . CFBDD is a validated method for inhibitor discovery and has been used to successfully identify inhibitors for a variety of proteins, including historically challenging targets such as KRAS G12C(31, 32).

In this study, we screened CHIKV nsP2pro against a 6,120-compound covalent fragment library using an enzymatic activity assay and discovered 153 hit compounds using a 50% inhibition threshold. Subsequent characterization of hits led to the identification of RA-0002034, a potent inhibitor of CHIKV nsP2pro activity and CHIKV replication. This covalent fragment, RA-0002034, contains a vinyl sulfone warhead, and exhibited a nanomolar IC_50_ in a biochemical protease activity assay. RA-0002034 showed specific covalent modification of the active site cysteine (C478) and minimal reactivity against a panel of cysteine proteases. Additionally, RA-0002034 inhibited replication of four alphaviruses in cell-based viral replication assays. Taken together, we have identified RA-0002034 as a chemical probe for CHIKV and as a promising starting point for further therapeutic development.

## Results

### Screening-compatible FRET-based Assay Development

To develop a platform for the identification of CHIKV nsP2pro inhibitors, we optimized an HTS-compatible FRET-based CHIKV nsP2pro activity assay, adapted from similar previously reported assays(33, 34). The FRET-based assay utilizes recombinantly expressed and purified CHIKV nsP2pro (Figure S1) and a 16-mer internally quenched fluorogenic peptide derived from the CHIKV nsP1/nsP2 cleavage site. As the peptide is cleaved by the protease, the Cy5 fluorophore is liberated from the quencher and fluorescence can be monitored (Figure 1A). Enzyme concentration, substrate concentration, and buffer conditions were optimized to give a robust signal to background ratio and suitable Z’ scores. We first determined the optimal buffer conditions by titrating components in a small design of experiment study, using constant nsP2 enzyme (1 μM) and substrate (10 μM) concentrations. The highest enzymatic activity was obtained in buffer containing 25 mM HEPES pH 7.2, 0.003% v/v Tween-20, and 1 mM DTT (Figure S2A-B). These conditions were used to titrate nsP2pro concentrations for an optimal signal window and determine the length of the linear range. We observed a robust signal at 150 nM nsP2pro, with Z’ scores >0.9 and a linear range of ∼2 hours (Figure S2C).

**Figure 1.**
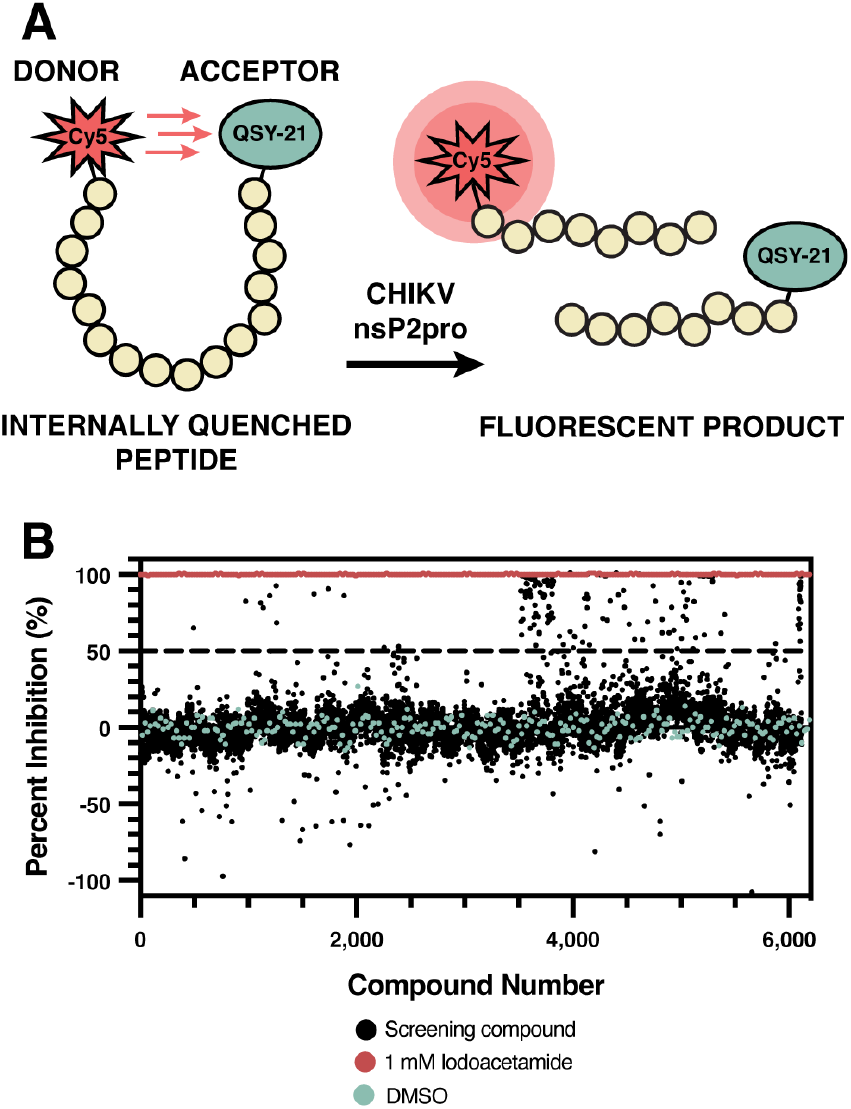
Covalent fragment screening. (A) Principle of HTS-compatible FRET-based assay; the internally quenched fluorescent peptide is cleaved in the presence of CHIKV nsP2pro to release fluorescent product. (B) Results of a 6,120-compound covalent fragment screen using the HTS-compatible FRET-based assay. A 50% inhibition threshold (dashed line) was used to define “active” compounds. Positive control (iodoacetamide) and negative control (DMSO) used for screening are shown in red and blue, respectively.

### Covalent Fragment Screening

Using the optimized HTS-compatible assay, we screened the Enamine Covalent Fragment Library (6,120-compounds) (Figure 1B). This library is composed of fragments tethered to acrylamide, chloroacetamide, fluorosulfonyl, nitrile, epoxide, aziridine, boronic acid, and vinyl sulfone warheads (Figure S3). The final compound concentration was 20 μM, with a final assay DMSO concentration of 2% v/v. Compounds were preincubated with CHIKV nsP2pro for 30 minutes, and the reaction with the internally quenched peptide was allowed to progress for 90 minutes before reading the fluorescence at 670 nm. To normalize the screening compound data, 1 mM iodoacetamide, which exhibits an IC_50_ of 15 μM (Figure S2D), was used as a positive control, and DMSO was used as a negative control. The average Z’ score of all plates was 0.92, well above the threshold of ∼0.5 to be considered a robust assay(35). Utilizing an activity threshold of 50% inhibition, 153 ‘actives’ were obtained, with an overall hit rate of 2.5% (Data Set 1 in Supporting Information). Of the 153 actives, 139 compounds contained either chloroacetamide or vinyl sulfone warheads. When adjusted for their overall representation in the library however, vinyl sulfone was the preferred warhead, as 39.5% of vinyl sulfones in the library showed greater than 50% inhibition (Table S1).

The 153 active compounds were clustered based on structural similarity using fragment-based descriptors (Figure S4). Using KNIME, Morgan fingerprints were generated for each compound, reduced to two dimensions using principal component analysis, and assigned into clusters using a k-means clustering algorithm. Representative compounds from each cluster, totaling 43 compounds, were repurchased (ChemSpace). DMSO stocks of repurchased compounds were dispensed in 10-point, 3-fold serial dilutions starting with a final concentration of 200 μM and tested using the previously described enzymatic activity assay. Of the 43 compounds tested, 20 compounds showed IC_50_s below 20 μM (Table S2, Figure S5-6). The 20 compounds with IC_50_s below 20 μM contained either an acrylamide, chloroacetamide, or vinyl sulfone warhead, with 15/20 compounds containing a vinyl sulfone warhead. The vinyl sulfone compounds were classified into two separate groups, internal or external vinyl sulfones, based on the location of the electrophile. Of the 43 compounds tested in dose-response format, RA-0002034 (Figure 2A) was the most potent (IC_50_ = 180 ± 20 nM) (Figure 2B).

**Figure 2.**
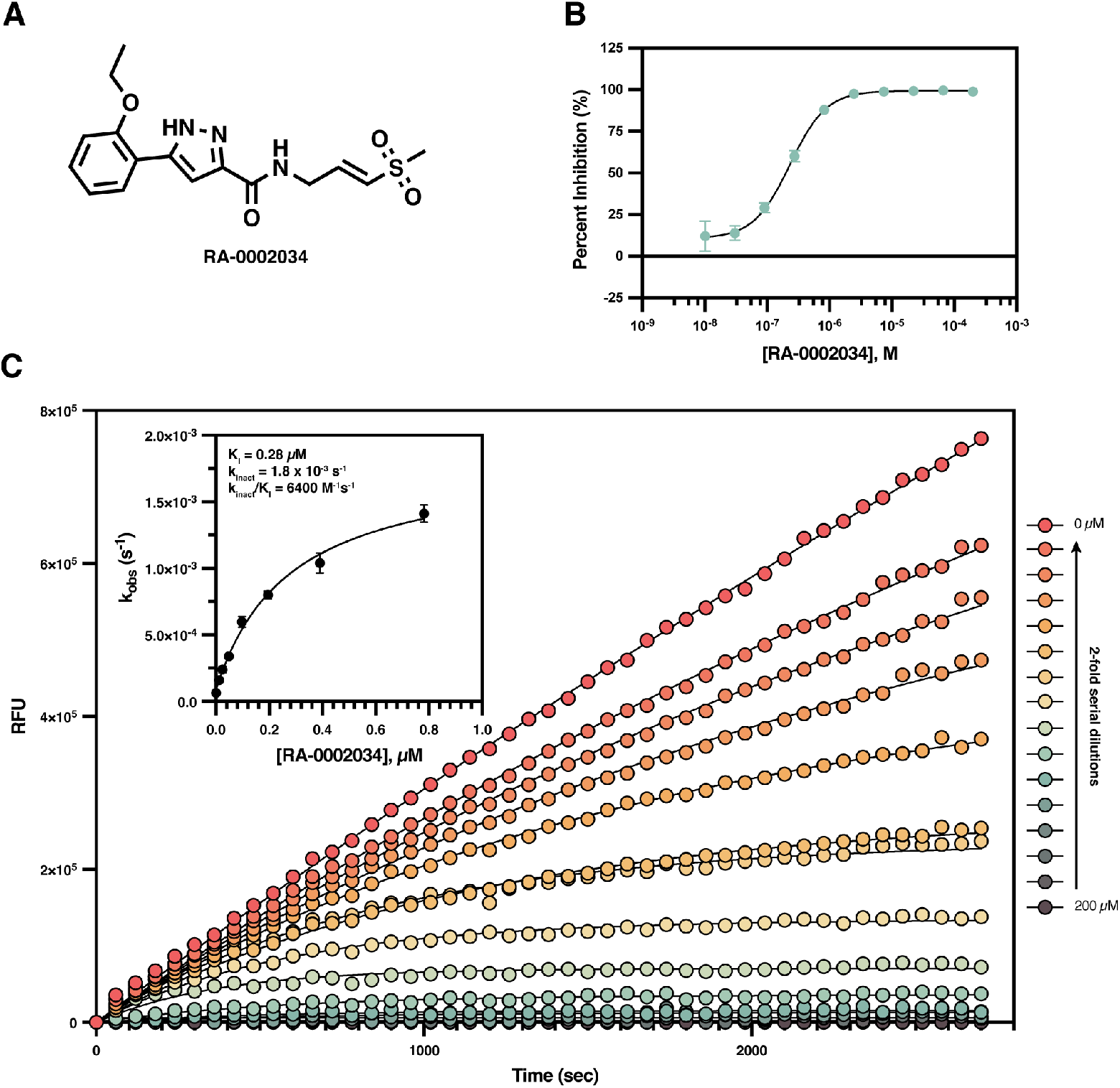
RA-0002034 potently inhibits CHIKV nsP2pro activity. (A) The structure of RA-0002034. (B). Dose-response curve of RA-0002034. 3-fold serial dilutions of RA-0002034 were prepared beginning from a starting concentration of 200 μM over 10 points and tested in the protease activity assay using the primary screening assay conditions. Data was fit in GraphPad Prism using a four-parameter Hill equation to yield an IC_50_ of 180 ± 20 nM. Data points represent mean ± standard deviation (n=3). (C) CHIKV nsP2pro is inhibited in a time-dependent manner by RA-0002034 as shown by a decrease in reaction rate (RFU/sec) over time. 2-fold serial dilutions of RA-0002034 were used to determine K_I_, k_inact_, and second order rate constant k_inact_/K_I_ (n=3).

### Evaluation of General Thiol Reactivity

One liability of covalent inhibitors is the propensity to react with off-target thiol-containing molecules. From the 20 compounds with IC_50_s less than 20 μM, 2-3 compounds from each warhead class were selected for reactivity analysis using a LC-MS-based GSH reactivity assay to determine a GSH t_1/2_ (Figure S7A). The compounds containing an external vinyl sulfone warhead were the most reactive with GSH, resulting in an average half-life of ∼5 min. The chloroacetamides and internal vinyl sulfones were modestly reactive, with half-lives ranging from 30 min to 4 hours. The most stable warheads were the acrylamides which were unreactive with GSH up to 24 hours (Figure S7B, Table 1). These results align with prior characterization of drug-like covalent fragments, which also found that external vinyl sulfones were rapidly reactive with free thiols whereas internal vinyl sulfones were relatively less reactive (36). As a secondary validation to the GSH results, an absorbance-based DTNB (Ellman’s reagent, 5,5’-dithiobis-2-nitrobenzoic acid) thiol reactivity assay was used to quantify thiol reaction rates of the fragment compounds (30). There was general rank-order agreement between the two assays, with the external vinyl sulfones remaining the fastest reacting of the warheads tested, with an average 1.5-fold increase in reactivity compared to chloroacetamides, a 60-fold increase in reactivity compared to the internal vinyl sulfones, and a ∼1000-fold increase compared to the acrylamides (Figure S8, Table S3). Due to the fast reactivity of external vinyl sulfones with thiols, the compounds containing that warhead were deemed promiscuously reactive and not pursued further (37).

**Table 1.** Experimental results for 20 compounds from the covalent fragment primary screen with IC_50_s below 20 μM in the enzymatic activity assay. Values shown in this table were obtained using the initial enzyme screening assay (before re-optimization of enzyme concentration to lower the tight binding limit; see Results and Figure S12).

| RA ID | Warhead | IC<sub>50</sub> (μM) | GSH t<sub>1/2</sub> (min) | K<sub>i</sub> (μM) | k<sub>inact</sub> (s<sup>-1</sup>) | k<sub>inact</sub>/K<sub>i</sub>(M<sup>-1</sup>s<sup>-1</sup>) |
| --- | --- | --- | --- | --- | --- | --- |
| RA-0002034 | Internal vinyl sulfone | 0.18 ± 0.02 | 130 | 0.28 ± 0.08 | 0.0018 ± 0.0002 | 6400 ± 990 |
| RA-0002352 | Internal vinyl sulfone | 0.39 ± 0.08 | | 2.2 ± 0.4 | 0.0024 ± 0.0003 | 1100 ± 60 |
| RA-0002482 | Internal vinyl sulfone | 0.59 ± 0.09 | | 5.0 ± 1.3 | 0.0023 ± 0.0003 | 460 ± 52 |
| RA-0002481 | Internal vinyl sulfone | 1.0 ± 0.2 | | 15 ± 2 | 0.0024 ± 0.0001 | 160 ± 20 |
| RA-0002056 | Internal vinyl sulfone | 1.2 ± 0.5 | 84 | 15 ± 1 | 0.0018 ± 0.0002 | 120 ± 6 |
| RA-0002505 | Internal vinyl sulfone | 1.2 ± 0.5 | | 19 ± 2 | 0.0021 ± 0.0002 | 110 ± 4 |
| RA-0002048 | External vinyl sulfone | 1.6 ± 0.2 | 7.2 | | | |
| RA-0002043 | Chloroacetamide | 1.7 ± 0.1 | 260 | | | |
| RA-0002054 | External vinyl sulfone | 3.4 ± 0.2 | 3.7 | | | |
| RA-0002039 | Chloroacetamide | 4.5 ± 2.3 | 72 | | | |
| RA-0002057 | Internal vinyl sulfone | 8.4 ± 1.5 | 200 | | | |
| RA-0002051 | External vinyl sulfone | 8.4 ± 0.3 | | | | |
| RA-0002058 | Acrylamide | 8.6 ± 2.8 | >1400 | | | |
| RA-0002055 | External vinyl sulfone | 9.1 ± 0.6 | | | | |
| RA-0002038 | External vinyl sulfone | 11 ± 2 | | | | |
| RA-0002037 | External vinyl sulfone | 12 ± 2 | | | | |
| RA-0002052 | External vinyl sulfone | 16 ± 2 | | | | |
| RA-0002047 | External vinyl sulfone | 17 ± 1 | | | | |
| RA-0002044 | Chloroacetamide | 18 ± 1 | 33 | | | |
| RA-0002035 | Acrylamide | 20 ± 7 | >1400 | | | |

### Time-Dependent Inhibition of CHIKV nsP2pro Activity

To validate the covalent mechanism of action of the inhibitory fragments, six compounds with IC_50_s below 1.5 μM, RA-0002034, RA-0002352, RA-0002482, RA-0002481, RA-0002056, and RA-0002505, were first evaluated in a time-course IC_50_ assay, with all six compounds containing an internal vinyl sulfone warhead. While a shift in IC_50_ over time was observed for all six compounds (Figure S9), this parameter does not fully describe inhibitory characteristics of covalent inhibitors. The second order rate constant k_inact_/K_I_ is the preferred method to characterize and rank the potency of covalent inhibitors for a specific target(38). k_inact_/K_I_ fully describes the transformation of enzyme to enzyme-inhibitor complex, taking into consideration the affinity of the reversible noncovalent interaction (K_I_) and the rate of covalent bond formation (k_inact_). To determine the k_inact_/K_I_ for the same set of six compounds, RA-0002034, RA-0002352, RA-0002482, RA-0002481, RA-0002056, and RA-0002505, we performed time-dependent enzyme inhibition experiments following the experimental setup outlined in Mons et al. (39). Briefly, the internally quenched peptide substrate was added to serially diluted compounds, and the reaction was initiated by the addition of CHIKV nsP2pro. The reaction progress was monitored immediately using a continuous read over 45 minutes. The time course plots were fit with Equation 1 to determine k_obs_ and replotted against inhibitor concentration to determine K_I_ and k_inact_ values for each inhibitor tested (Figure S10, Table 1). Notably, the k_inact_ values for all six compounds tested were within the margin of error from one another, however the K_I_ values varied, suggesting that the increases in potency are driven by an increased binding affinity, not an increase in reactivity.

### RA-0002034 Demonstrates the Most Promising Potency and Reactivity Profile

RA-0002034 initially yielded the lowest IC_50_ (180 ± 20 nM) using the primary screening assay conditions. RA-0002034 also showed modest reactivity in the GSH assay with a half-life of 130 min and low reactivity in the DTNB assay. RA-0002034 exhibited a K_I_ of 0.28 ± 0.08 μM with a k_inact_ of 1.8 x 10-3 ± 2.0 x 10^-4^ s^-1^, resulting in a k_inact_/K_I_ value of 6.4 x 10^3^ ± 9.9 x 10^2^ M^-1^s^-1^, 5-6-fold greater than the next closest compound (Figure 2C, Table 1). Based on the initial potency and reactivity characterization, RA-0002034, which contains a methyl vinyl sulfone warhead, became the top compound of interest. RA-0002034 was resynthesized (Scheme S1-3) and confirmed to be >95% pure and of the correct structure as measured by NMR, HPLC, and HRMS. (Figure S11A-H). The synthesized product obtained was solely the (*E*) isomer, whereas the initial commercial stock was a mixture of the (*E*) and (*Z*) isomers. During synthesis, it was observed that RA-0002034 could undergo an intermolecular aza-Michael reaction to form a cyclic 6-member ring, however synthesis conditions were identified to obtain pure acyclic product (Scheme S1-3) and to avoid the inactive cyclic product (Figure S13B).

Due to the IC_50_ of RA-0002034 (180 ± 20 nM) approaching the concentration of CHIKV nsP2pro used in the dose-response experiments (150 nM; i.e. the ‘tight binding limit’), the enzymatic assay was reoptimized to reduce the concentration of CHIKV nsP2pro. CHIKV nsP2pro titrations and time-course experiments showed 20 nM CHIKV nsP2pro was sufficient to produce a robust fluorescent signal (Z’ > 0.6) at a 90-minute endpoint, and the signal remained in the linear range (Figure S12). The activity of resynthesized RA-0002034 was tested in the re-optimized dose-response activity assay using 20 nM CHIKV nsP2pro and was found to have an IC_50_ of 58 ± 17 nM (Figure S13A), comparable to the inhibition seen with the repurchased compound (180 ± 20 nM). The resynthesized compound was used for all subsequent experiments.

### RA-0002034 Covalent Modification is Specific for the Catalytic Cysteine

To confirm our hypothesis that RA-0002034 inhibits CHIKV nsP2pro by alkylating the active site cysteine, mass spectrometry was used to determine which cysteine or cysteines formed an adduct with RA-0002034. CHIKV nsP2pro domain contains six cysteines, four of which are solvent exposed (32; PDB ID: 4ZTB). After incubation with a serial dilution of RA-0002034 or DMSO for 30 minutes and tryptic digestion, proteomic analysis by LC-MS/MS revealed a modification corresponding to the mass shift (+349.11 Da) of RA-0002034 on the A475-K481 (ANVCWAK) tryptic peptide (Figure 3A). This modified peptide was observed in all RA-0002034-treated samples and absent in a DMSO-only control (Figure 3B). The modification was confidently localized to the catalytic cysteine (C478). The intensity of the RA-0002034 modified peptide was also dose-dependent, corresponding with the amount of RA-0002034 added to the sample (Figure 3C). No modifications of other cysteine-containing tryptic peptides were observed. These results strongly indicate that RA-0002034 is indeed active site-directed and is specific for only C478, the catalytic cysteine of CHIKV nsP2pro.

**Figure 3.**
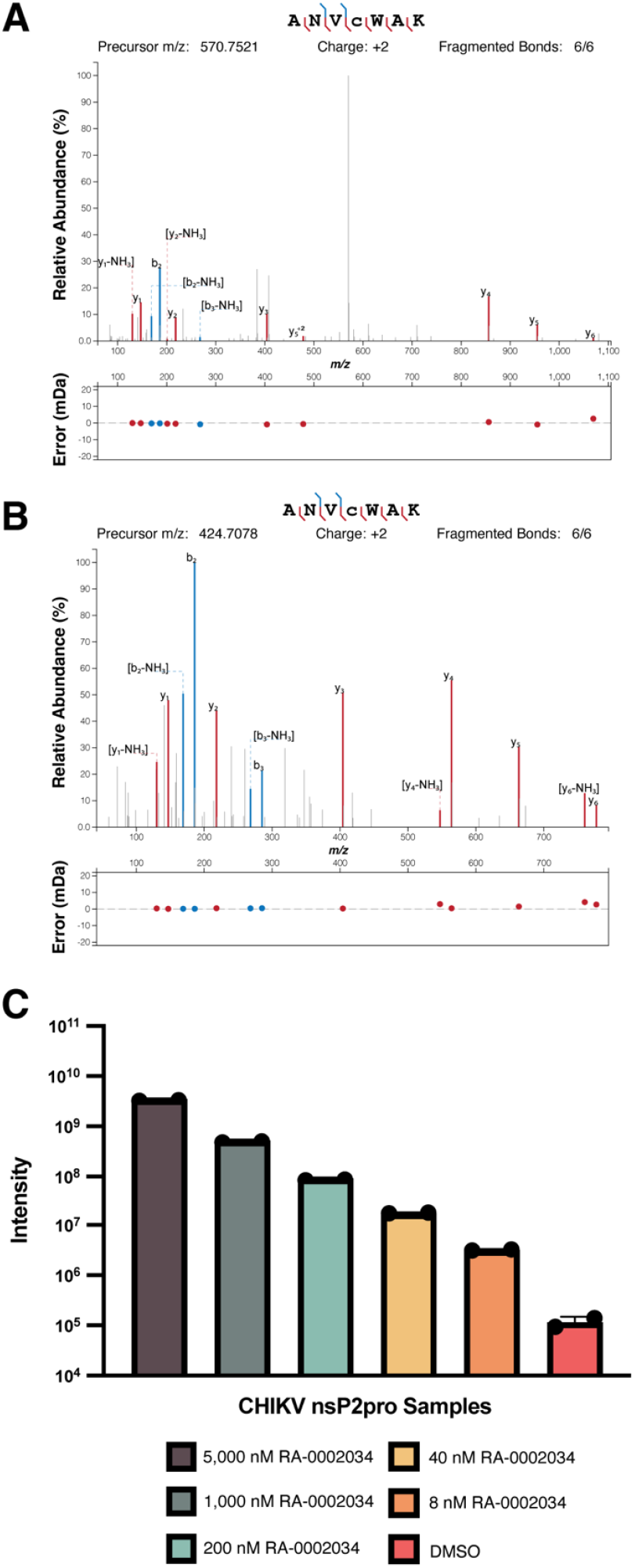
RA-0002034 selectively reacts with catalytic cysteine. Trypsin digestion product peptide spectra containing the catalytic cysteine (ANVCWAK) from samples incubated with (A) 5 μM RA-0002034 or (B) DMSO. (C) The intensity of peaks corresponding with C478-RA-0002034 adduct is dose dependent.

### RA-0002034 is Selective for CHIKV nsP2pro

One drawback to TCIs is potential toxicity due to off-target reactivity. Undesired inhibition of human cysteine proteases (calpains, cathepsins, etc.) and subsequent downstream pharmacological effects are a potential concern with an irreversible cysteine protease inhibitor. To determine whether RA-0002034 exhibits off-target protease reactivity, we screened a panel of 13 diverse cysteine proteases, comprised of caspase 1, calpain 1, calpain 2, cathepsins B, C, H, K, L, S, V, papain, SARS-CoV-2 Mpro, and TEV protease. RA-0002034 exhibited an IC_50_ value greater than 100 μM against all cysteine proteases tested, apart from cathepsin S, for which RA-0002034 had an IC_50_ of 40 μM, ∼700-fold less potent compared to CHIKV nsP2pro, suggesting protease inhibition by RA-0002034 is selective for nsP2pro (Figure S14, Table S4).

### Modeling Predicts Binding of RA-0002034 into S1’-S3 Subsites

To predict the binding pose of RA-0002034 in the CHIKV nsP2pro active site, RA-0002034 was covalently docked into CHIKV nsP2pro. In its predicted pose, the compound resides in the substrate binding groove, occupying the S1’-S3 subsites (Figure 4A-B)(33). In the S2 subsite, the oxygen of the ethoxyphenyl group on RA-0002034 is predicted to hydrogen bond with the indole nitrogen of W549. In the S3 subsite, the RA-0002034 pyrazole is predicted to have hydrogen bonding interactions with the Y544 hydroxyl and the backbone amide nitrogen of N547. The amide nitrogen of RA-0002034 is predicted to have hydrogen bonding interactions with the backbone carbonyl of N547 (Figure 4C).

**Figure 4.**
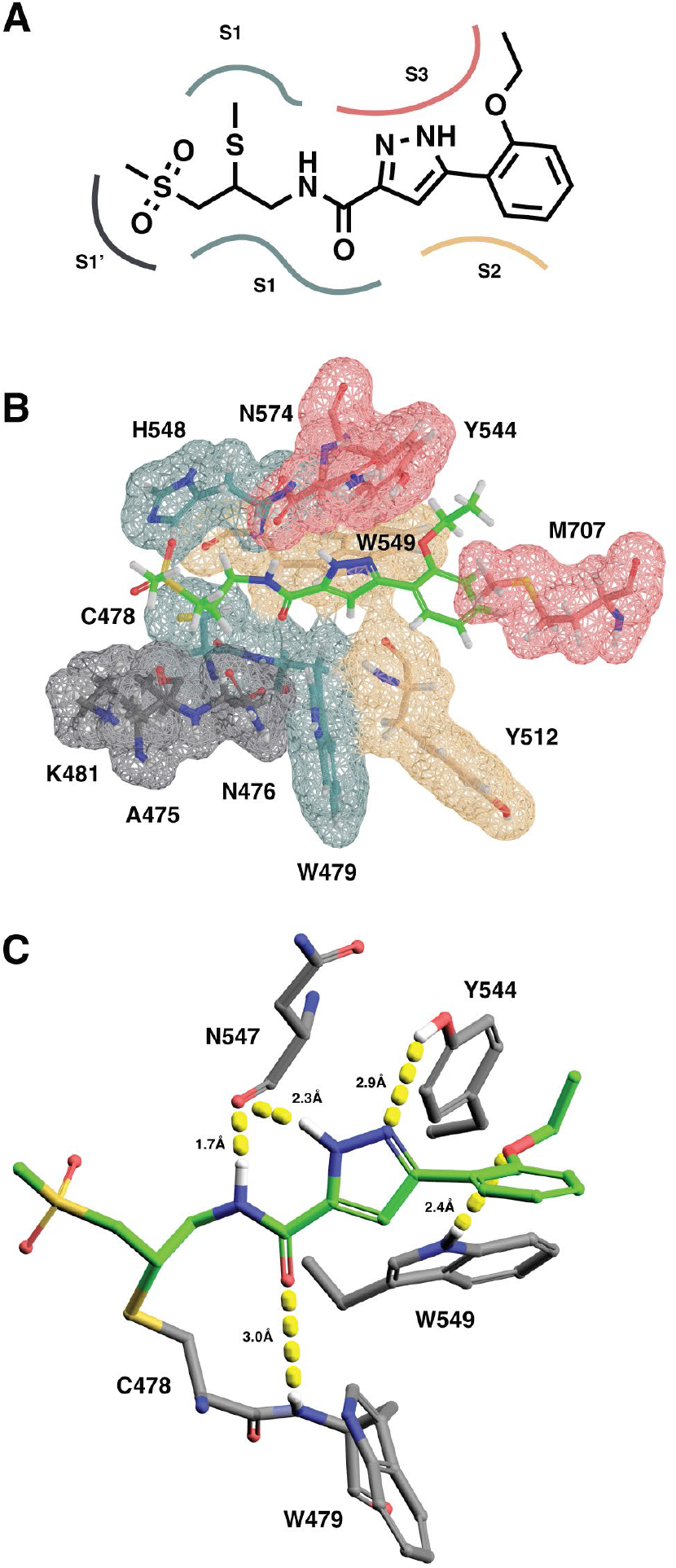
RA-0002034 was covalently docked into CHIKV nsP2pro (PDB 3TRK). (A) Diagram of substrate recognition subsites and (B) surface view of RA-0002034 docked in substrate binding cleft. S1’, S1, S2, and S3 subsites are colored grey, blue, yellow, and red, respectively. (C) Intermolecular interaction between RA-0002034 (green) and CHIKV nsP2pro residues (grey).

To study flexibility of the nsP2pro active site we performed all-atom Molecular Dynamics (MD) simulations. A high-resolution X-ray structure (PDB ID: 3TRK) was used as the initial structure for our simulations. We found that the loop containing the catalytic histidine (N547-H548-W549) is the region of the protein with the highest conformational flexibility (Figure S15). In VEEV crystal structures, the conformation of this loop changes when in complex with the peptidomimetic inhibitor E-64d. Specifically, VEEV His546 flips in orientation towards the catalytic cysteine, the flexibility of which may aid in substrate recognition and selectivity(40),(41). In the docked pose, RA-0002034 is positioned directly beneath this flexible loop. Because of the flexibility of the active site, intermolecular interactions may occur between RA-0002034 and CHIKV nsP2pro that are not visible by docking RA-0002034 to a rigid crystal structure. Confirmation of key residues that drive the binding interaction with RA-0002034 will require future co-crystallography and/or resistance mutation analysis.

### RA-0002034 Inhibits Viral Replication

Encouraged by the potency and selectivity of RA-0002034 for CHIKV nsP2pro, we investigated the inhibition of CHIKV replication by RA-0002034 utilizing a nanoluciferase (nLuc) reporter assay. The CHIKV nLuc reporter assay uses an attenuated CHIKV vaccine strain (181/25)(42) with a nLuc reporter inserted downstream of the capsid gene, and luciferase activity was used as a measure of CHIKV replication and polyprotein expression (Figure 5A). MRC5 cells were pretreated for 1 hour with a 4-fold serial dilution of RA-0002034 starting at 10 μM and assessed for viral replication at 6 hours post-infection (hpi). In parallel, we also used a VEEV TC83 nLuc reporter, which contains nLuc inserted into the nsP3 gene, to determine the alphavirus specificity of RA-0002034 (Figure 5B). RA-0002034 inhibited CHIKV replication with an EC_50_ of 11 ± 4 nM (Figure 5C) and VEEV replication with an EC_50_ of 320 ± 21 nM (Figure 5D), suggesting activity across multiple alphaviruses. Cell viability was also assessed at 24 hpi employing ATP production as a marker of cell health using Cell Titer Glo (CTG) (Promega). No toxicity was observed at the highest RA-0002034 concentration of 10 μM in the CTG assay (Figure 5E). Additional hit compounds from the primary screen were also tested in the nLuc and CTG assays (Figure S16-18, Table S5). However, RA-0002034 exhibited the most potent inhibition in the CHIKV reporter antiviral assay, correlating well with the inhibition observed in the enzymatic assay.

**Figure 5.**
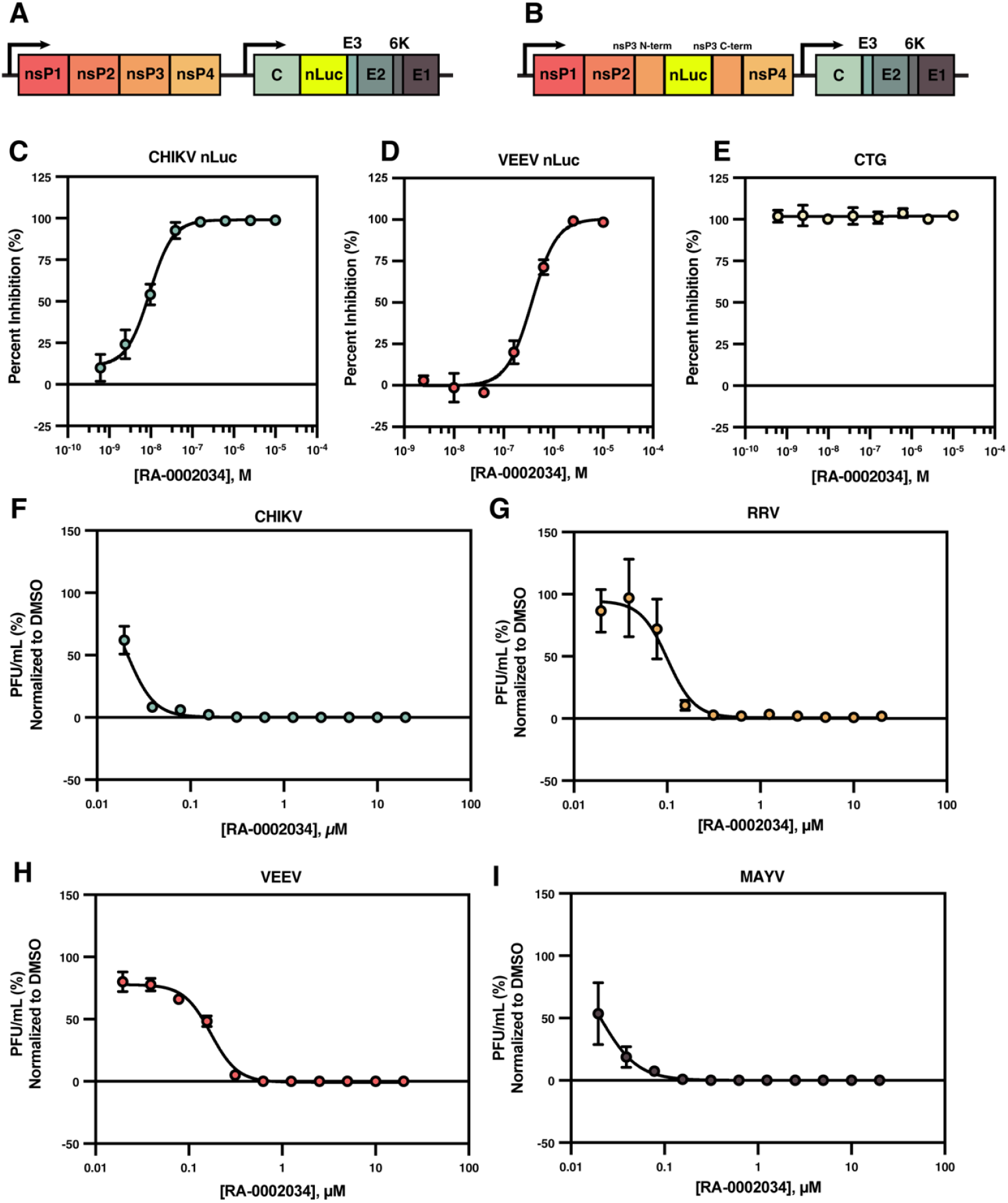
Cell-based antiviral potency of RA-0002034. Schematic of (A) CHIKV nLuc and (B) VEEV nLuc reporter assays. MRC5 cells were pretreated with a serial dilution of RA-0002034 and infected with either CHIKV nLuc (C) or VEEV nLuc (D). CTG assay was used to measure cell viability (E). Plots contain mean with standard deviation (n=3). CHIKV (F), RRV (G), VEEV (H), and MAYV (I) were tested in plaque formation assays using a serial dilution of RA-0002034. Plots contain mean with standard deviation (n=3).

To further validate and test the spectrum of RA-0002034 antiviral activity, we performed plaque formation assays over a serial dilution of RA-0002034 to measure reduction in viral titer against CHIKV, VEEV, MAYV, and RRV. RA-0002034 effectively reduced viral titers of all four viruses. RA-0002034 was too potent against CHIKV and MAYV to confidently fit for EC_90_ values, as both had ∼50% inhibition of plaque formation at the lowest dose of 19.5 nM, with reportable EC_90_s of <40 nM. EC_90_s were calculated for RRV and VEEV, 200 ± 41 nM and 360 ± 32 nM, respectively (Figure 5F-I), which together demonstrate that RA-0002034 is able to effectively inhibit viral replication across a broad spectrum of alphaviruses.

### Development of a Negative Control Chemical Probe

While RA-0002034 is promising for progression to hit-to-lead in an early drug discovery pipeline, the compound also has utility for the study of CHIKV as a chemical probe. Recently, quality criteria for covalent chemical probes have been established by Hartung et al. (37). A covalent chemical probe must have a measured and reported k_inact_/K_I_, an assessment of intrinsic chemical reactivity (GSH t_1/2_ > 5 min), cellular activity below 1 μM, a primary site of covalent interaction, and a selectivity factor over identified off-targets. RA-0002034 satisfies all the requirements for a high-quality covalent chemical probe; with a k_inact_/K_I_ of 6.4 x 10^3^ M^-1^s^-1^, a GSH t_1/2_ of 130 minutes, a sub-μM cellular potency of 11 nM, an identified modification site of C478, and selectivity over off-target cysteine proteases.

A valuable addition to a covalent chemical probe is the availability of a negative control probe that has reduced on-target potency while retaining the electrophilic warhead. To develop a negative control probe, RA-0003161 was synthesized (Scheme S4, Figure S19A-B). RA-0003161 retains the same core structure as RA-0002034, however, RA-0003161 contains a methoxymethyl substituent on the core pyrazole (Figure S20A). In the biochemical protease activity assay, RA-0003161 inhibited nsP2 protease with an IC_50_ of 30 ± 7 μM, a ∼500-fold decrease in potency relative to RA-0002034 (Figure S20B). RA-0003161 was also far less potent in the cell-based CHIKV and VEEV nLuc reporter assays, with EC_50_s of 3 μM and >10 μM, respectively; a ∼300-fold and >30-fold decrease in antiviral activity in the CHIKV and VEEV nLuc reporter assays, respectively, relative to RA-0002034 (Figure S20C-D). Both RA-0002034 and RA-0003161 showed no cell toxicity at 10 μM (Figure S20E).

## Discussion

CHIKV and other related alphavirus cause debilitating disease across the globe (2). There is currently a lack of high-quality, validated small molecule probes in the chemical toolbox for the study of alphaviruses as well as a lack of inhibitors of alphavirus replication. In this study, we utilized a covalent fragment screening approach to discover the covalent chemical probe RA-0002034, which potently inhibits CHIKV replication through inhibition of CHIKV nsP2pro by specific covalent modification of the catalytic cysteine residue. RA-0002034 has a k_inact_/K_I_ of 6.4 x 10^3^ ± 9.9 x 10^2^ M^-1^s^-1^ and a cell-based EC_50_ of 11± 4 nM in a CHIKV nLuc reporter assay.

RA-0002034 contains a methyl vinyl sulfone warhead, and interestingly, Zhang et al., reported an inhibitor of VEEV nsP2pro activity and VEEV replication, referred to as compound 11, which contains similar methyl vinyl sulfone(19). While compound 11 exhibits μM potency against VEEV replication in HeLa (EC_50_ = 1.6 μM) and Vero E6 (EC_50_ = 2.4 μM) cells, RA-0002034 potently inhibits both VEEV and CHIKV replication, with EC_50_s of 320 ± 21 nM and 11 ± 4 nM, respectively in nLuc reporter assays in MRC5 cells. Compound 11 has not yet been reported to inhibit viruses other than VEEV. Despite these apparent differences, the two independent discoveries of vinyl-sulfone-containing inhibitors of alphavirus nsP2 proteases reinforces the potential of this chemotype for inhibitor development of nsP2pro function and alphavirus replication. Vinyl sulfone inhibitors have also shown utility as proteosome inhibitors(43) and inhibitors of parasitic proteases(44). The *Trypanosoma cruzi* inhibitor K777 (or K11777) targets the *T. cruzi* cysteine protease cruzipain with a phenyl vinyl sulfone warhead. K777 effectively treats *T. cruzi* infection in cell, mouse, and dog models with no apparent toxicity (45–47), again showing promise for this warhead-containing chemotype.

One concern with covalent inhibitors is irreversible adduct formation with off-target proteins. To evaluate potential off-target inhibition, RA-0002034 was tested against a panel of cysteine proteases. RA-0002034 did not inhibit any of the cysteine proteases tested, except for cathepsin S which was weakly inhibited with an IC_50_ of 40 μM, well above the IC_50_ for CHIKV nsP2pro (58 ± 17 nM). Additionally, no reduction in cellular viability was observed in a CTG assay. In future studies, development of an activity-based probe for proteomic profiling of RA-0002034 cellular targets will be necessary for understanding the full scope of RA-0002034 selectivity.

While LC-MS/MS studies have validated that RA-0002034 specifically modifies the catalytic cysteine of CHIKV nsP2pro, how the compound interacts with the surrounding active site sub-pockets has not been fully elucidated. Docking and MD studies indicate that RA-0002034 most likely binds to the nsP2pro active site within the S1’-S3 pockets, placing the vinyl sulfone warhead in close proximity to the nucleophilic catalytic cysteine. In the docked pose, the RA-0002034 pyrazole resides directly beneath the flexible loop containing the active site cysteine. As part of RA-0002034 probe development, we synthesized RA-0003161, which contains a methoxymethyl moiety attached to the pyrazole, resulting in a significant drop in potency compared to RA-0002034 in both enzymatic assays as well as antiviral reporter assays. This result makes sense in the context of the docking model, as a bulky substituent added to the pyrazole would cause steric clashes with the flexible loop near the active site. A co-crystal structure of RA-0002034 covalently linked to CHIKV nsP2pro will provide a full understanding of which CHIKV nsP2pro active site residues make key intermolecular contacts with RA-0002034 to drive binding and electrophile alignment with the active site cysteine (C478).

We find that RA-0002034 effectively inhibits replication of both Old World (CHIKV, RRV) and New World (VEEV, MAYV) alphaviruses, suggesting the important potential of broad-spectrum alphavirus efficacy. While RA-0002034 potently inhibits each of the alphaviruses tested in the titer reduction panel, RA-0002034 was the least potent against VEEV (EC_90_ = 360 ± 32 nM). One potential explanation for the modest reduction in potency against VEEV compared to CHIKV is the difference in the nsP2pro active site architecture between the viruses. In the docked pose Y544, which is located in the S3 subsite, is in close enough proximity to a RA-0002034 pyrazole nitrogen to form a hydrogen bond. CHIKV Y544 is conserved amongst CHIKV, MAYV, and RRV; however, in VEEV an isoleucine resides in that position. The loss of a hydrogen bond interaction between the Y544 hydroxyl and the pyrazole nitrogen could be responsible for the reduction in potency seen in VEEV reporter and titer reduction assays. Further evaluation of specific intermolecular interactions will provide insight into compound features needed for broad-spectrum activity.

Here, we report the discovery of the covalent chemical probe RA-0002034, the most potent CHIKV nsP2pro inhibitor to date, which exhibits a profile suitable for advancement to hit-to-lead studies. RA-0002034 effectively inhibits CHIKV nsP2pro enzymatic activity, viral replication of CHIKV and VEEV in nLuc reporter assays, and CHIKV, VEEV, MAYV, and RRV replication in titer reduction assays. Proteomic analysis demonstrated that the covalent reaction is specific for the CHIKV nsP2pro catalytic cysteine. Additionally, RA-0002034 exhibits excellent selectivity for CHIKV nsP2pro, with limited to no off-target inhibition in a panel of diverse cysteine proteases. Taken together, RA-0002034 represents a promising starting point towards the development of a CHIKV and/or broad-spectrum alphavirus therapeutic by targeted inhibition of nsP2pro.

## Materials and Methods

### Protein production

A bacterial expression vector containing 6xHis-tagged CHIKV nsP2pro was designed based on the existing crystal structure (PDB 4ZTB) (AA1009-1329 [UniProt accession Q5XXP4]). CHIKV nsP2pro protein was expressed in Rosetta2 cells and clarified cell lysates were purified using Ni-NTA resin (Qiagen) to produce highly purified and active enzyme. CHIKV nsP2pro (1009-1329) was used in all subsequent experiments. The vector was transformed into Rosetta2 cells and cell cultures were grown at 37 ºC to an OD_600_ of 0.6 and protein expression was induced using 200 μM IPTG and grown overnight at 15 ºC. Cell pellets were resuspended in Buffer A (50 mM NaHPO4 pH 7.8, 500 mM NaCl, 10 mM imidazole, 20% glycerol, 1 mM β-mercaptoethanol) containing 1 mg/mL lysozyme and Roche cOmplete protease inhibitor (without EDTA) and sonicated with a 20 second pulse at 40% amplitude and 40 second rest for 12 cycles. The cell lysate was clarified by centrifugation at 12,000 RPM at 4 ºC for 60 minutes. 1 mL of washed Ni-NTA agarose resin was added to clarified lysate and incubated at 4 ºC for 4 hours on an orbiter. The resin was isolated by decanting into a 5 mL gravity column. The column was washed with 5 mL Buffer A then washed with 2 mL 10% Buffer B (50 mM NaHPO4 pH 7.8, 500 mM NaCl, 250 mM imidazole, 20% glycerol, 1 mM β-mercaptoethanol) to 90% Buffer A. Protein was eluted with 100% Buffer B and dialyzed overnight using Slide-a-lyzer (ThermoFisher) cassettes in dialysis buffer (25 mM HEPES pH 7.5, 500 mM NaCl, 20% Glycerol, 10 mM DTT). Protein concentration was measured using a NanoDrop spectrophotometer, flash-frozen in liquid nitrogen, and stored at -80 ºC.

### CHIKV nsP2pro activity assays

An internally quenched peptide derived from the CHIKV 1/2 cleavage site (QLEDRAGA/GIIETPRG) was designed with an N-terminal QSY®-21 quencher and a C-terminal Cy5 fluorophore (Lifetein) for use as a substrate in enzymatic cleavage assays. 10 μL enzymatic reactions were performed in 384-well, black, non-binding plates (Corning 3820). For the primary screening campaign with the Enamine Covalent Fragment Library, a Mosquito liquid handler (STP LabTech) was used to transfer 200 nL from mother plates containing 1 mM compound in DMSO into assay-ready plates for a final screening concentration of 20 μM. For dose-response plate preparation, three-fold compound dilutions were prepared in DMSO using a Tecan Evo and transferred to assay-ready plates using the Mosquito liquid handler as above. For the assay, a stock solution of 10X CHIKV nsP2pro activity buffer (250 mM HEPES pH 7.2) was filtered through a 0.2 μm filter and stored at 4 ºC. Prior to running the assay, a 2x solution of nsP2pro (300 nM) was prepared in 2x activity buffer (50 mM HEPES pH 7.2, 0.006% Tween20, 2 mM DTT) and 2x fluorogenic substrate peptide (20 μM) was prepared in reagent-grade water. 5 μL of 2x enzyme was dispensed into assay ready plates containing 200 nL compound and allowed to pre-incubate with compound for 30 minutes at room temperature with slow shaking. To initiate the reaction, 5 μL of 2x peptide substrate solution was added and incubated for 1.5 hours at room temperature. Reagents were dispensed into assay-ready plates using a Combi Multidrop (Thermo Scientific). Fluorescence was measured at 685 nm using a PerkinElmer Envision 2105 plate reader. For time-dependent enzyme inhibition experiments, solutions and plates were prepared as previously described, however 5 μL of peptide was dispensed into the plate prior to the addition of 5 μL enzyme, rapidly shaken for 5s and then read continuously on the PerkinElmer Envision 2105 plate reader for 2 hours. Raw data was normalized to background fluorescence and fit to Equation 1 to yield k_obs_ values. For primary screening and dose-response experiments, data were uploaded into the UNC Center for Integrative Chemical Biology and Drug Discovery (CICBDD) database (ScreenAble Solutions) for storage and analysis.

### GSH reactivity assay

100 μL reactions were prepared containing 2.5 mM GSH, 500 μM covalent compound, and 800 μM Rhodamine B in 100 mM KHPO_4_, 10% acetonitrile (ACN). The final mixture was injected onto an LC-MS at 30–60-minute intervals until depletion of compound reached >50%. The AUC of covalent compound peaks were calculated and normalized to the internal standard by dividing the AUC compound by AUC Rhodamine B. The second order reaction was considered a pseudo first-order reaction due to [GSH] >> [Compound], therefore reaction half-life was determined as t_1/2_=ln_2_/k, where k was the rate of depletion of compound. k was calculated by using a linear regression of the natural logs of the normalized AUC time course measurements.

### DTNB thiol reactivity assay

A DTNB solution containing 50 μM DTNB (ThermoFisher) and 200 μM TCEP (Hampton Research) was prepared in 20 mM sodium phosphate pH 7.4, 150 mM NaCl and incubated for 5 minutes at room temperature to produce TNB^2-^. 50 μL of the TNB^2-^ solution was transferred to each well of a 384-well clear plate (Griener Bio-One #781162) containing 2.5 μL of electrophilic fragment (10 mM in DMSO, final concentration 500 μM). The plate was rapidly shaken, sealed with an optically clear plate seal, and transferred to a PerkinElmer Envision 2105 plate reader, acquiring absorbance measurements at 412 nm every 5 minutes for 16 hours at 37 ºC. Once complete, the background absorbance of samples was normalized to an unreduced DTNB control to correct for compound absorbance in the 400 nm wavelength range. Concentrations of TNB^2-^ and electrophilic fragments were calculated at each time point and fit to a second-order reaction, making the value of the rate constant k equal to the slope of ln([A][B0]/[B][A0]). [A0] and[B0] are the initial concentrations of the electrophilic fragment (500 μM) and TNB^2-^ (100 μM), and [A] and [B] are remaining concentrations at each time point calculated from the absorbance measurements. A linear regression of the linear range of measurements was used to determine k.

### Sample preparation for modification site mapping

CHIKV nsP2 pro (1.5 mg/mL) was incubated with various concentrations of RA-0002034 (5 μM-0.008μM) for 30 minutes at room temperature in 25 mM HEPES pH 7.2 and 1 mM DTT. After incubation, excess compound was removed using a spin concentrator. Protein concentration was normalized to 1 mg/mL, flash frozen, and stored at -80 ºC. 20 μg of the sample was diluted to 50 μL with 8M urea, reduced with 5 mM DTT and alkylated with 15 mM iodoacetamide. Samples were diluted to reduce the concentration of urea under 1M and then digested using 0.5 μg/μl trypsin overnight at 37 ºC, then desalted the next day using C18 spin columns (Pierce).

### LC-MS/MS analysis

The samples were analyzed by LC/MS/MS using an Easy nLC 1200 coupled to a QExactive HF mass spectrometer (Thermo Scientific). Samples were injected onto an Easy Spray PepMap C18 column (75 μm id × 25 cm, 2 μm particle size) (Thermo Scientific) and separated over a 120 min method. The gradient for separation consisted of 5–38% mobile phase B at a 250 nL/min flow rate, where mobile phase A was 0.1% formic acid in water and mobile phase B consisted of 0.1% formic acid in 80% ACN. The QExactive HF was operated in data-dependent mode where the 15 most intense precursors were selected for subsequent fragmentation. Resolution for the precursor scan (m/z 350–1600) was set to 60,000 with a target value of 3 × 10^6^ ions. MS/MS scans resolution was set to 15,000 with a target value of 1 × 10^5^ ions, 100 ms injection time. The normalized collision energy was set to 27% for HCD. Dynamic exclusion was set to 30 s, peptide match was set to preferred, and precursors with unknown charge or a charge state of 1 and ≥ 8 were excluded.

### LC-MS/MS data analysis

Raw data were processed using Proteome Discoverer (Thermo Scientific, version 2.5). For the in-vitro analysis, data were searched against a reviewed Uniprot *E*.*coli* BL21 database (containing 4,156 sequences; downloaded April 2023), appended with the CHIKV-nsp2 sequence and a common contaminants database (MaxQuant, 245 sequences), using the Sequest HT search algorithm within Proteome Discoverer. Enzyme specificity was set to trypsin, up to two missed cleavage sites were allowed; RA-0002034 (+349.11 Da) and carbamidomethylation of Cys, as well as oxidation of Met were set as variable modifications. A precursor mass tolerance of 10 ppm and fragment mass tolerance of 0.02 Da were used. Label-free quantification (LFQ) was enabled. Peptides were filtered based on a 1% false discovery rate (FDR). Statistical analysis was performed in Perseus (version 1.6.14.0) using the peptide-level LFQ intensities (unimputed). Student’s t-test (two-tailed, uncorrected) was performed for the RA-0002034 to DMSO comparison and a p-value < 0.01 was considered statistically significant. LFQ log2 fold change ratios were calculated and an absolute log2 ratio ±-1.5 was considered significant.

### Luciferase Reporter Assays

MRC5 fibroblasts cultured in DMEM supplemented with 10% FBS and L-glutamine were treated with an eight-point four-fold serial dilution of compound or DMSO one hour prior to infection with nLuc reporter virus for CHIKV (181/25-nLuc-Capsid) or VEEV (TC83-nLuc-nsP3) at an MOI of 0.1 or left uninfected. After 6 hours, cells infected with virus were treated with Nano-Glo® luciferase assay reagent to detect nLuc luminescence. To assess cell viability, uninfected cells were treated with CellTiterGlo® 2.0 reagent (Promega). In each case, luminescence was measured on Promega GloMax® GloMax plate reader. Percent inhibition and viability were calculated by normalization to DMSO control wells. EC_50_ and CC_50_ values were calculated from compounds run in technical and biological triplicate using a three parameter Hill equation.

### Alphavirus Titer Reduction Assay

10 mM compound stocks were used to prepare 10-point dose response curves ranging from 40 μM to 0.156 μM by diluting compounds 1:1 with DMEM supplemented with 5% FBS and 1% PSG. Media only and DMSO controls (5 μL/mL) were included. One hour before infection, diluted compounds and controls were added to 48-well plates of confluent normal human dermal fibroblasts in triplicate. Cells were infected with a multiplicity of infection equal to 0.5-2 depending upon the virus used in the assay. At 2 hpi, infection media was removed, wells were washed two times with PBS, and fresh media containing compound was added to cells. At 24-48 hpi, 20 μL samples of supernatant were transferred to 96-well plates for titering by limiting dilution plaque assays to determine levels of virus production. For the plaque assay, 10-fold serial dilutions were made in the 96-well plates and 100 μL of each dilution was transferred into wells of confluent monolayers of Vero cells in a 48-well plate. At 2 hpi, the cells were overlaid with 250 μL of carboxymethyl cellulose prepared in DMEM supplemented with 5% FBS and PSG. At 48 hpi, the plates were fixed with 3.7% formalin and stained with 0.4% methylene blue dye. Plaques were counted using a stereomicroscope. Data was tabulated in Excel and prepared for EC_90_ calculation by normalization of all data to DMSO controls. EC_50_ and EC_90_ values were calculated using a nonlinear regression equation with Prism v7 by GraphPad Software, Inc.

### Docking

The docking of RA-0002034 was performed using Glide Covalent Docking protocol(48). The structure of the CHIKV nsP2pro (PDB ID:3TRK) was prepared using the Schrödinger protein preparation wizard(49, 50), which provides minor structure optimization, ensuring the best performance of the following Glide runs. We used the standard setting of the wizard: add and optimize the positions of the hydrogen atoms and alter residue ionization/tautomer states using PROPKA(51, 52) at pH 7, remove water molecules further than 3 Å away from the HET atoms, and perform restrained structure minimization with the OPLS4 force field(53) converging the heavy atoms to RMSD of 0.3 Å. The prepared structure was used to generate the Glide receptor grid. The center of the grid box was placed on the CYS residue with the inner and outer box sizes equal to 20 Å and 40 Å, respectively. The structure of RA-0002034 was prepared using LigPrep tool to ensure that we sample all possible conformational models of the ligand during the docking protocol. All generated states of the ligand were covalently docked into the receptor grid prepared in the previous steps. We used Glide CovDock protocol for the docking and scoring. The best Glide models were selected based on the Glide docking score. The top scored pose was selected as a structural model of the complex.

## Acknowledgments

The authors thank Dr. Arunima Sikdar for reviewing the experimental data and Peter Buttery for assistance in establishing the LC-MS-based GSH assay.

## Figures and Tables

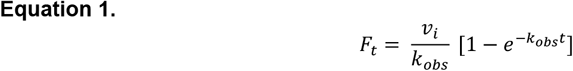

