## Supplementary Information for "Discovery of a cell-active chikungunya virus nsP2 protease inhibitor using a covalent fragment-based screening approach"

**This PDF file includes:**

**Supplemental Figures S1 to S21**
**Supplemental Tables S1 to S5**
**Supplemental Schemes S1 to S5**
**Supplemental Methods**
**Supplemental References**

**Other supplementary materials for this manuscript include the following:**

**Dataset S1**

**Supporting Information**


| Table of Contents | Page |
|---|---|
| Figure S1. Purification and activity of CHIKV nsP2pro. | S2 |
| Figure S2. Optimization of HTS-compatible enzymatic assay. | S3 |
| Figure S3. Enamine covalent fragment library information. | S4 |
| Figure S4. Clustering of active compounds from covalent fragment screen. | S5 |
| Table S1. Warhead distribution in library and actives. | S6 |
| Table S2. Dose-response validation of 43 repurchased active compounds. | S7 |
| Figure S5. Dose-response validation of 43 repurchased active compounds. | S8 |
| Figure S6. Chemical structures of 20 compounds $IC_{50}$ < 20 μM | S12 |
| Figure S7. GSH reactivity of active compounds. | S13 |
| Figure S8. DTNB thiol reactivity. | S14 |
| Table S3. Thiol reactivity of covalent fragments. | S14 |
| Figure S9. Time-dependent $IC_{50}$ experiments. | S15 |
| Figure S10. Time-dependent inhibition for compounds 18, 10, 16, 21, and 15. | S16 |
| Scheme S1. Synthesis of CHIKV nsP2pro covalent hit RA-0002034 (75). | S17 |
| Scheme S2. Synthesis of 5-(2-ethoxyphenyl)-1H-pyrazole-3-carboxylic acid (S1). | S17 |
| Scheme S3. Synthesis of (E)-3-(methylsulfonyl)prop-2-en-1-amine hydrochloride (S2). | S18 |
| Figure S11A-B. $^1$H and $^{13}$C NMR of intermediate S1. | S19 |
| Figure S11C-D. $^1$H and $^{13}$C NMR of intermediate S2. | S20 |
| Figure S11E-F. $^1$H and $^{13}$C NMR of RA-0002034 (75). | S21 |
| Figure S11G. LC-MS and HPLC purity of RA-0002034 (75). | S22 |
| Figure S11H. HRMS spectrum of RA-0002034 (75). | S23 |
| Figure S12. CHIKV nsP2pro enzymatic assay re-optimization. | S24 |
| Figure S13. Dose-response validation of resynthesized RA-0002034 and cyclic RA-0002442. | S25 |
| Figure S14. Cysteine protease panel. | S26 |
| Table S4. Cysteine protease panel. | S27 |
| Figure S15. CHIKV nsP2pro MD simulations | S28 |
| Figure S16. Covalent fragment CHIKV nLuc dose-response results. | S29 |
| Figure S17. Covalent fragment VEEV nLuc dose-response results. | S30 |
| Figure S18. CTG cell viability. | S31 |
| Table S5. nLuc reporter assay results for covalent fragments. | S32 |
| Scheme S4. Synthesis of RA-0003161 (154). | S33 |
| Figure S19A-B. $^1$H and $^{13}$C NMR of RA-0003161 | S34 |
| Scheme S5. Synthesis of methoxymethyl (MOM)-protected pyrazole carboxylic acid (**S3**) | S35 |
| Figure S20A-B. $^1$H and $^{13}$C NMR of compound S3 | S36 |
| Figure S21. RA-0003161 inhibition data. | S37 |
| Supplemental Methods | S38 |
| Supplemental References | S40 |

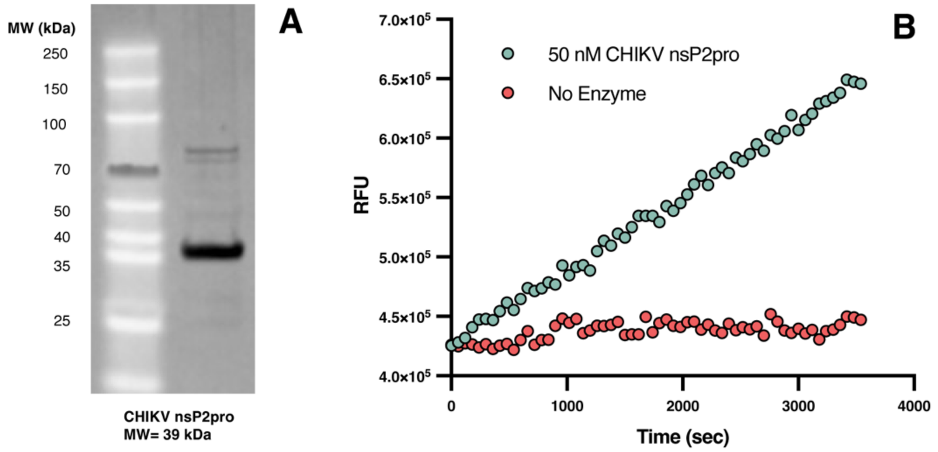


**Figure S1. Purification and activity of CHIKV nsP2pro.** (A) SDS-PAGE gel of purified CHIKV nsP2pro. (B) CHIKV nsP2pro purified enzyme is active and cleaves internally quenched peptide.

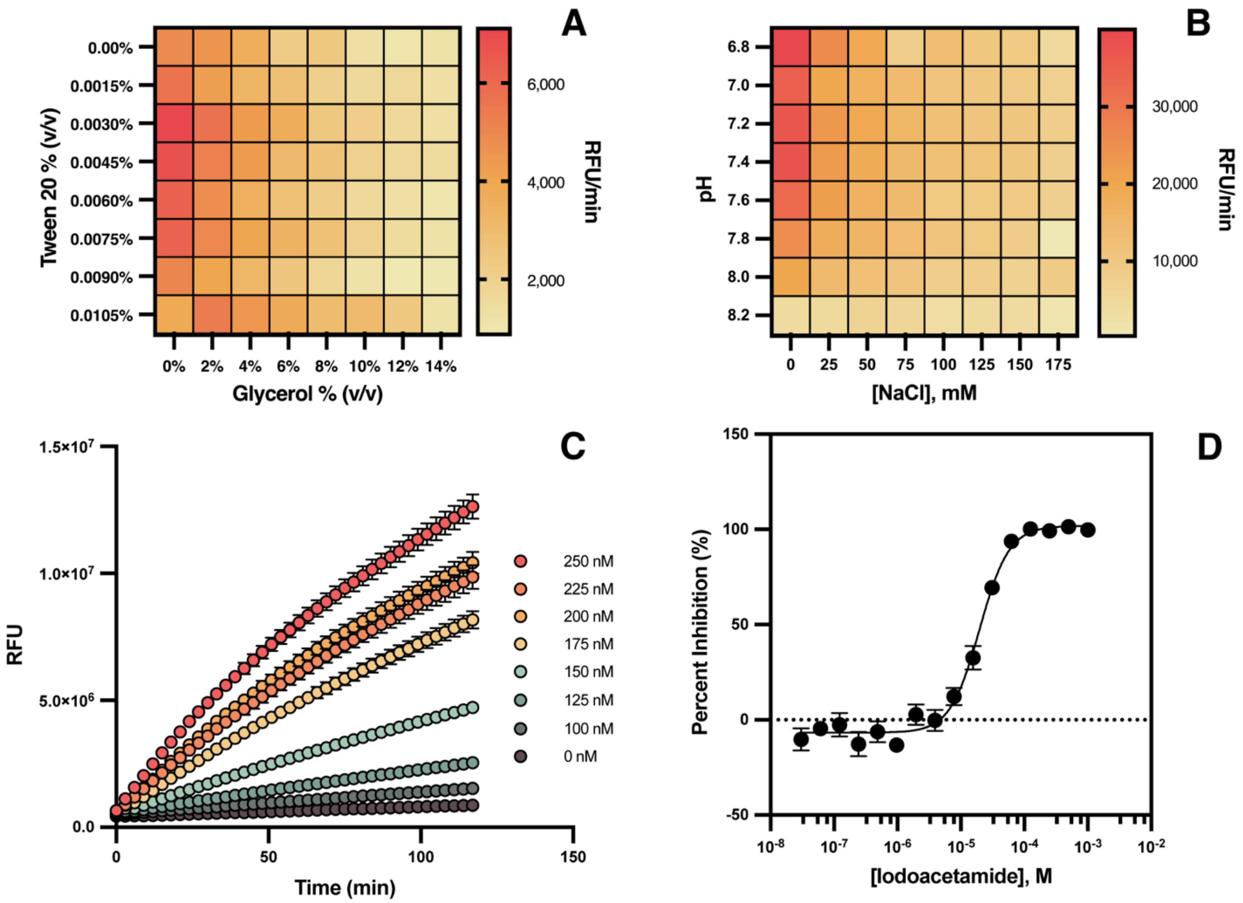


**Figure S2. Optimization of HTS-compatible enzymatic assay.** (A-B) Buffer components are titrated in 2D format. (C) Optimized buffer conditions were used to titrate enzyme concentration. (D) Using optimized enzyme and buffer concentrations, iodoacetamide was tested in dose response format and determined to have an $IC_{50}$ of 20 μM (n=3, data points represent mean ± standard deviation).

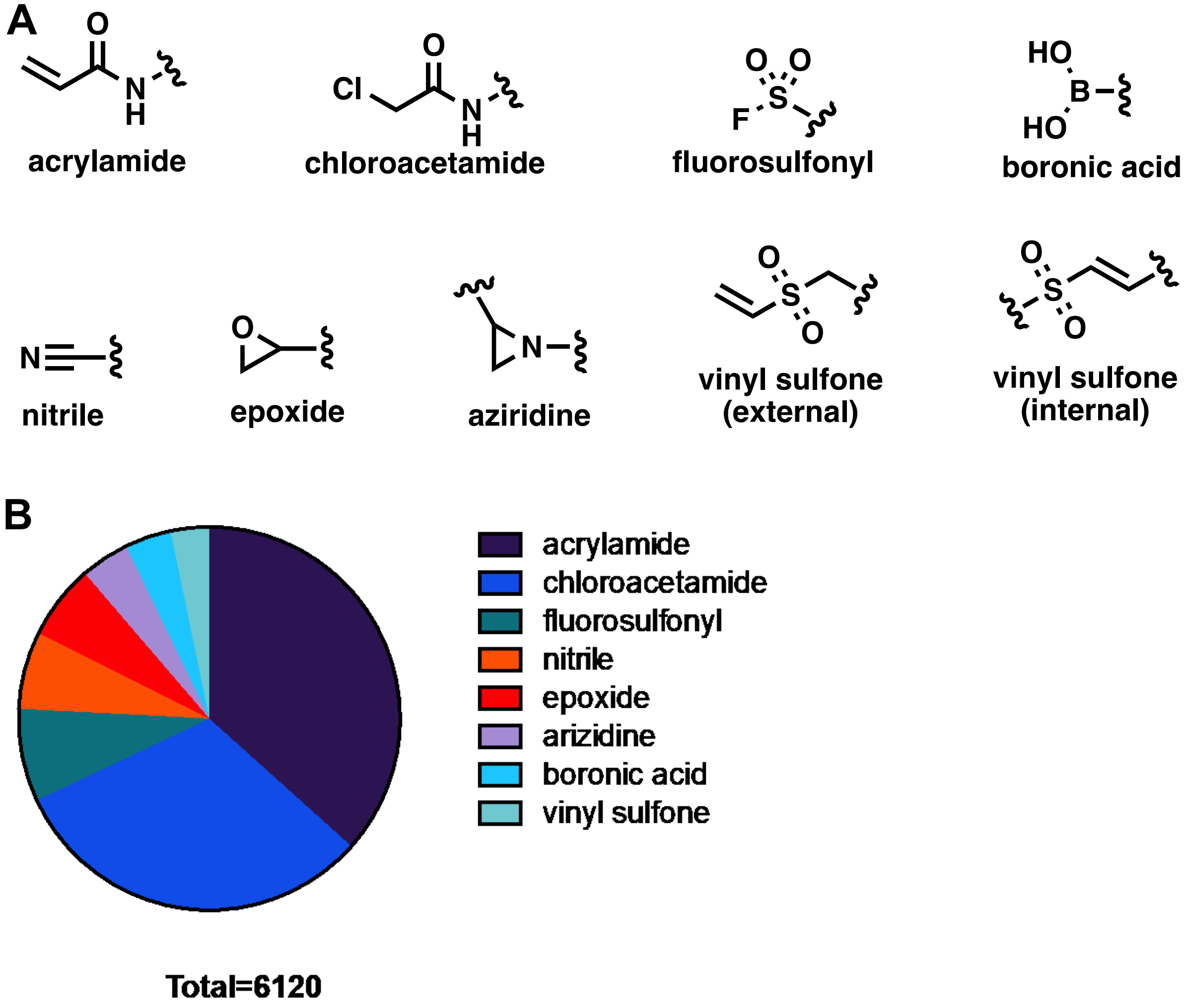


**Figure S3. Enamine covalent fragment library information.** (A) Structures of warheads present in the library. (B) Distribution of warheads in the library.

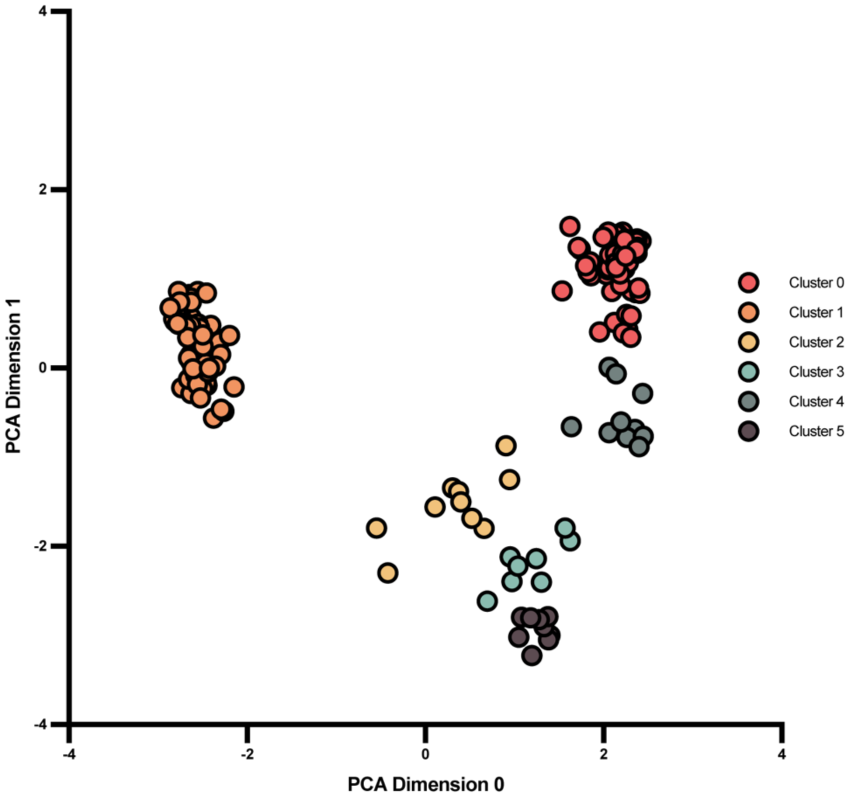


**Figure S4. Clustering of active compounds from covalent fragment screen.** Morgan fingerprints were generated for 153 active compounds and reduced to 2D using PCA. Clusters were assigned using k-means clustering algorithm (n=6).

**Table S1.** Warhead distribution in library and actives.

| Warhead | # in library | # active | Percent Active |
|---|---|---|---|
| acrylamide | 2240 | 12 | 0.54% |
| chloroacetamide | 1920 | 60 | 3.13% |
| fluorosulfonyl | 480 | 0 | 0.00% |
| nitrile | 400 | 1 | 0.25% |
| epoxide | 387 | 0 | 0.00% |
| aziridine | 253 | 0 | 0.00% |
| boronic acid | 240 | 1 | 0.42% |
| vinyl sulfone | 200 | 79 | 39.50% |
| Total | 6,120 | 153 | 2.50% |

**Table S2.** Dose-response validation of 43 repurchased active compounds.

| Compound | RA ID | Warhead | $IC_{50}$ (μM) |
|---|---|---|---|
| 75 | RA-0002034 | Internal vinyl sulfone | 0.20 ± 0.06 |
| 18 | RA-0002352 | Internal vinyl sulfone | 0.39 ± 0.08 |
| 10 | RA-0002482 | Internal vinyl sulfone | 0.59 ± 0.09 |
| 16 | RA-0002481 | Internal vinyl sulfone | 1.0 ± 0.2 |
| 21 | RA-0002056 | Internal vinyl sulfone | 1.2 ± 0.5 |
| 15 | RA-0002505 | Internal vinyl sulfone | 1.2 ± 0.5 |
| 11 | RA-0002048 | External vinyl sulfone | 1.6 ± 0.2 |
| 46 | RA-0002043 | Chloroacetamide | 1.7 ± 0.1 |
| 29 | RA-0002054 | External vinyl sulfone | 3.4 ± 0.2 |
| 4 | RA-0002039 | Chloroacetamide | 4.5 ± 2.3 |
| 56 | RA-0002057 | Internal vinyl sulfone | 8.4 ± 1.5 |
| 30 | RA-0002051 | External vinyl sulfone | 8.4 ± 0.3 |
| 70 | RA-0002058 | Acrylamide | 8.6 ± 2.8 |
| 66 | RA-0002055 | External vinyl sulfone | 9.1 ± 0.6 |
| 51 | RA-0002038 | External vinyl sulfone | 11 ± 2 |
| 107 | RA-0002037 | External vinyl sulfone | 12 ± 2 |
| 124 | RA-0002052 | External vinyl sulfone | 16 ± 2 |
| 99 | RA-0002047 | External vinyl sulfone | 17 ± 1 |
| 83 | RA-0002044 | Chloroacetamide | 18 ± 1 |
| 52 | RA-0002035 | Acrylamide | 20 ± 7 |
| 147 | RA-0002046 | External vinyl sulfone | 22 ± 4 |
| 13 | RA-0002480 | Internal vinyl sulfone | 23 ± 2 |
| 120 | RA-0002053 | External vinyl sulfone | 23 ± 2 |
| 118 | RA-0002036 | External vinyl sulfone | 24 ± 4 |
| 132 | RA-0002050 | External vinyl sulfone | 24 ± 3 |
| 128 | RA-0002049 | Internal vinyl sulfone | 30 ± 5 |
| 123 | RA-0002060 | Acrylamide | 33 ± 5 |
| 135 | RA-0002045 | Chloroacetamide | 41 ± 17 |
| 153 | RA-0002059 | Acrylamide | 41 ± 24 |
| 139 | RA-0002040 | Chloroacetamide | 49 ± 7 |
| 115 | RA-0002042 | Chloroacetamide | 65 ± 50 |
| 140 | RA-0002041 | Chloroacetamide | 74 ± 9 |
| 117 | RA-0002477 | Acrylamide | 130 ± 68 |
| 71 | RA-0002476 | Acrylamide | 140 ± 111 |
| 88 | RA-0002473 | Acrylamide | 140 ± 52 |
| 85 | RA-0002474 | Acrylamide | >200 |
| 95 | RA-0002475 | Acrylamide | >200 |
| 62 | RA-0002478 | Acrylamide | >200 |
| 148 | RA-0002479 | Nitrile | >200 |
| 9 | RA-0002483 | Internal vinyl sulfone | >200 |
| 144 | RA-0002484 | Boronic acid | >200 |
| 149 | RA-0002506 | Nitrile | >200 |
| 36 | RA-0002507 | Chloroacetamide | >200 |

**Figure S5.** Dose-response validation of 43 repurchased active compounds.

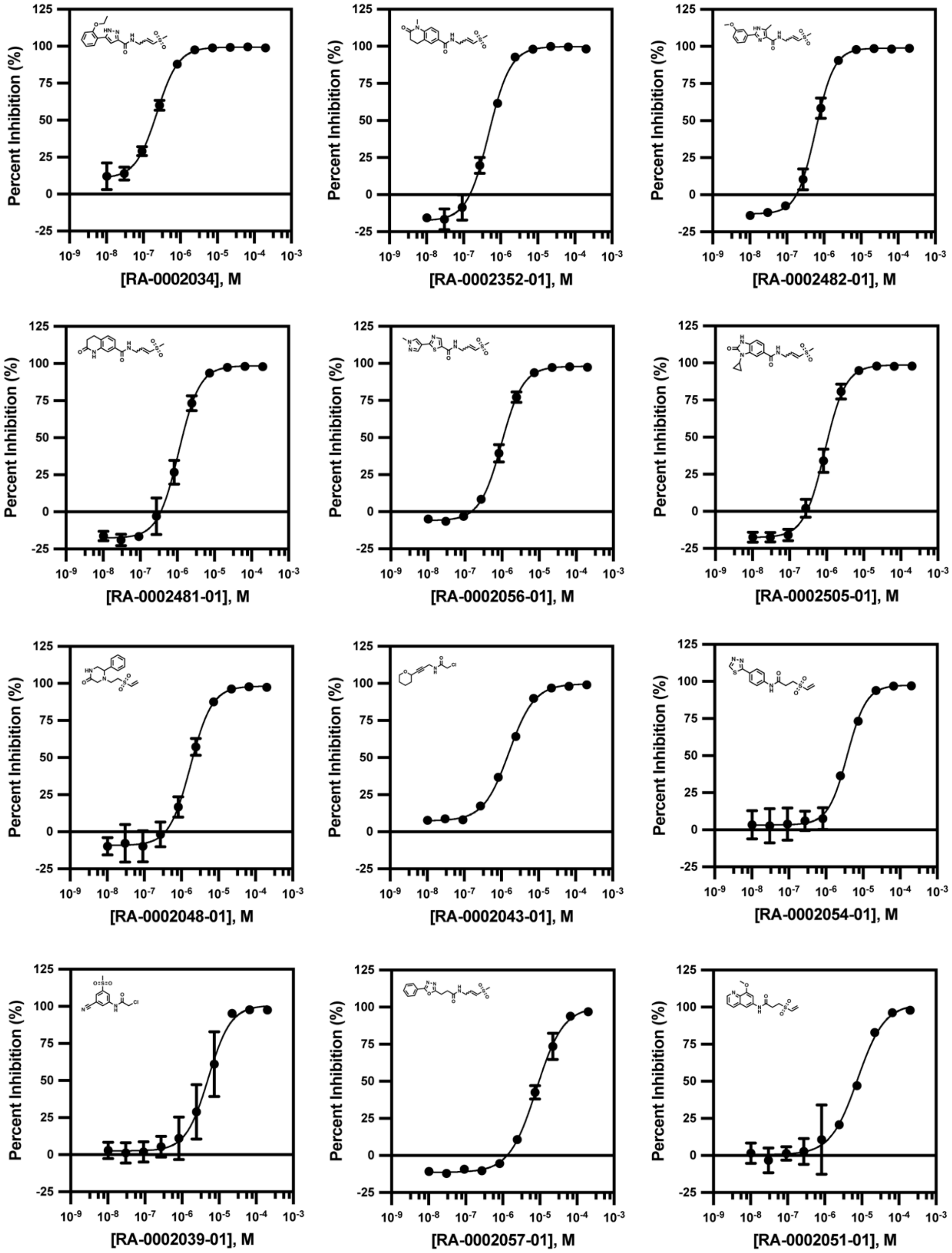

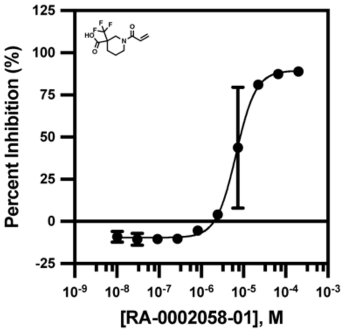
Percent Inhibition (%)
[RA-0002058-01], M
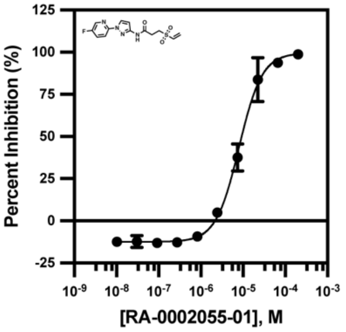
Percent Inhibition (%)
[RA-0002055-01], M
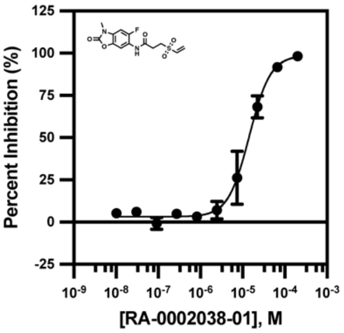
Percent Inhibition (%)
[RA-0002038-01], M
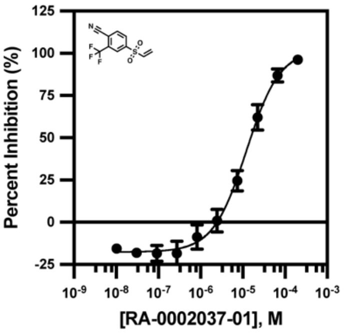
Percent Inhibition (%)
[RA-0002037-01], M
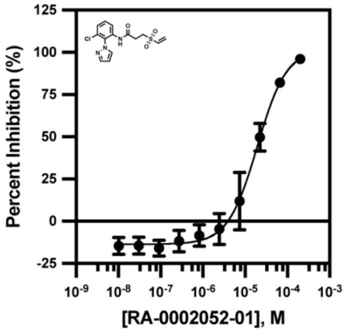
Percent Inhibition (%)
[RA-0002052-01], M
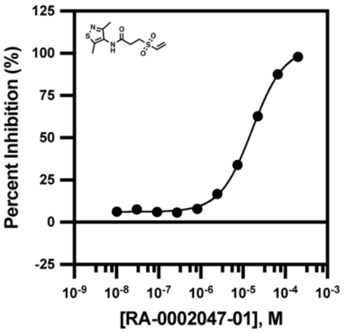
Percent Inhibition (%)
[RA-0002047-01], M
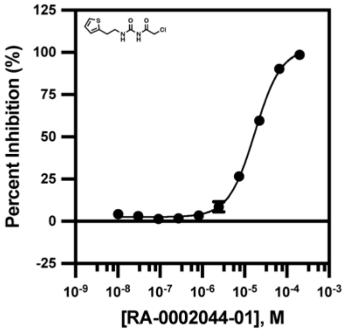
Percent Inhibition (%)
[RA-0002044-01], M
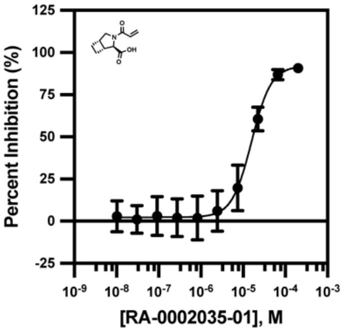
Percent Inhibition (%)
[RA-0002035-01], M
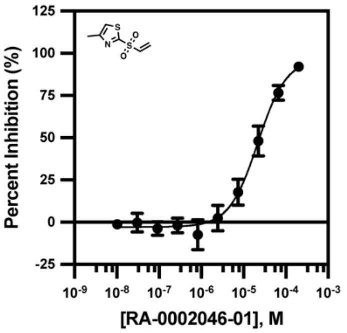
Percent Inhibition (%)
[RA-0002046-01], M
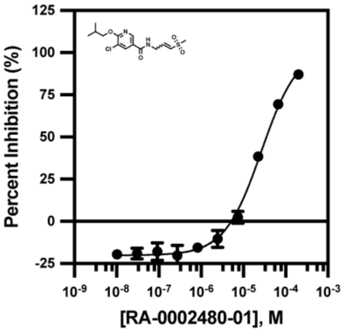
Percent Inhibition (%)
[RA-0002480-01], M
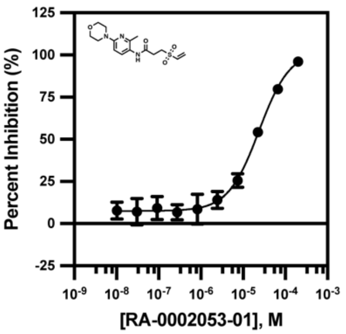
Percent Inhibition (%)
[RA-0002053-01], M
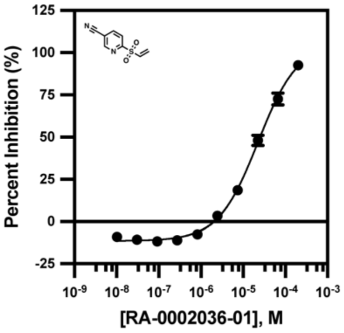
Percent Inhibition (%)
[RA-0002036-01], M

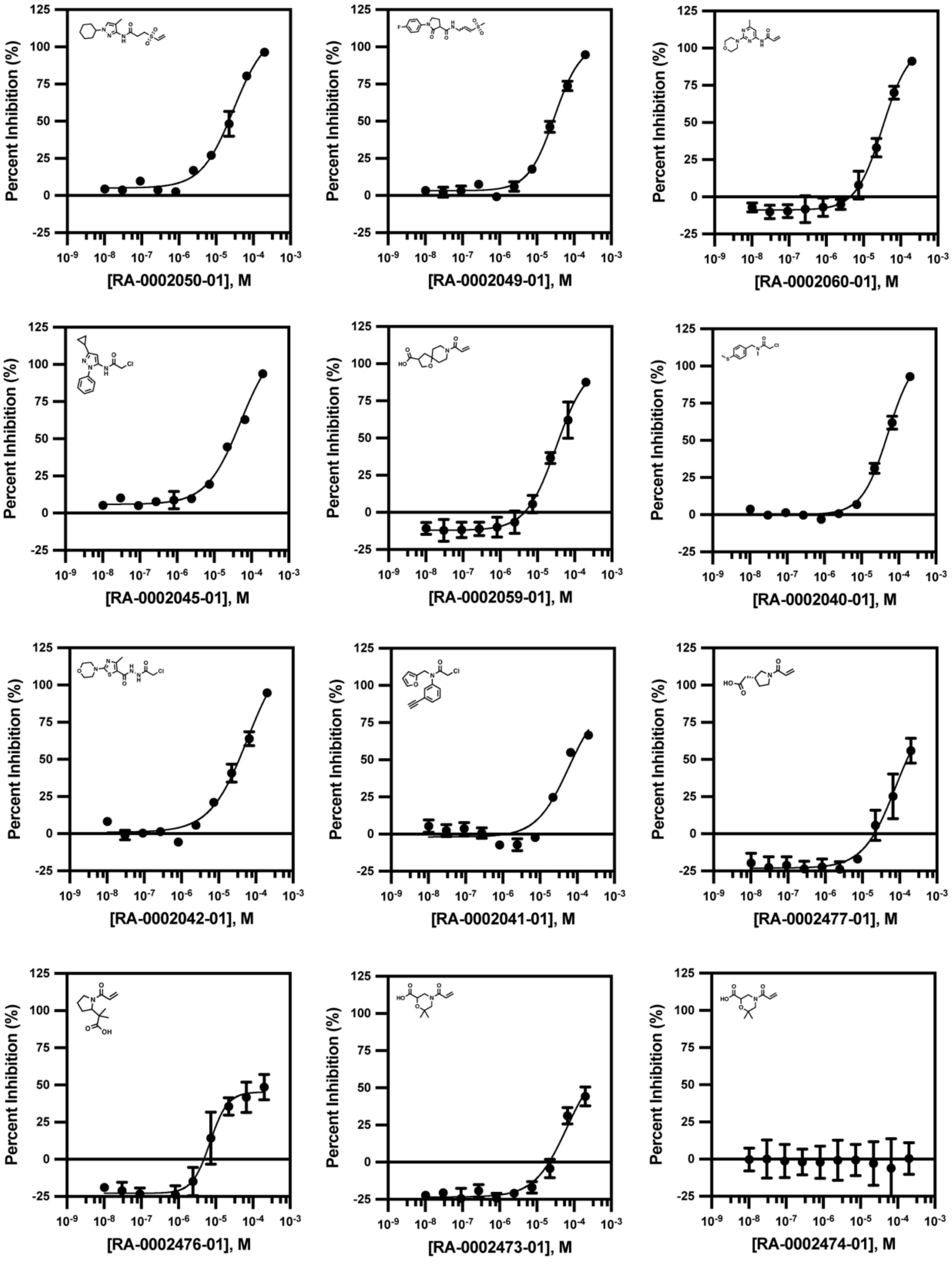
Percent Inhibition (%)
[RA-0002050-01], M
[RA-0002049-01], M
[RA-0002060-01], M
[RA-0002045-01], M
[RA-0002059-01], M
[RA-0002040-01], M
[RA-0002042-01], M
[RA-0002041-01], M
[RA-0002477-01], M
[RA-0002476-01], M
[RA-0002473-01], M
[RA-0002474-01], M

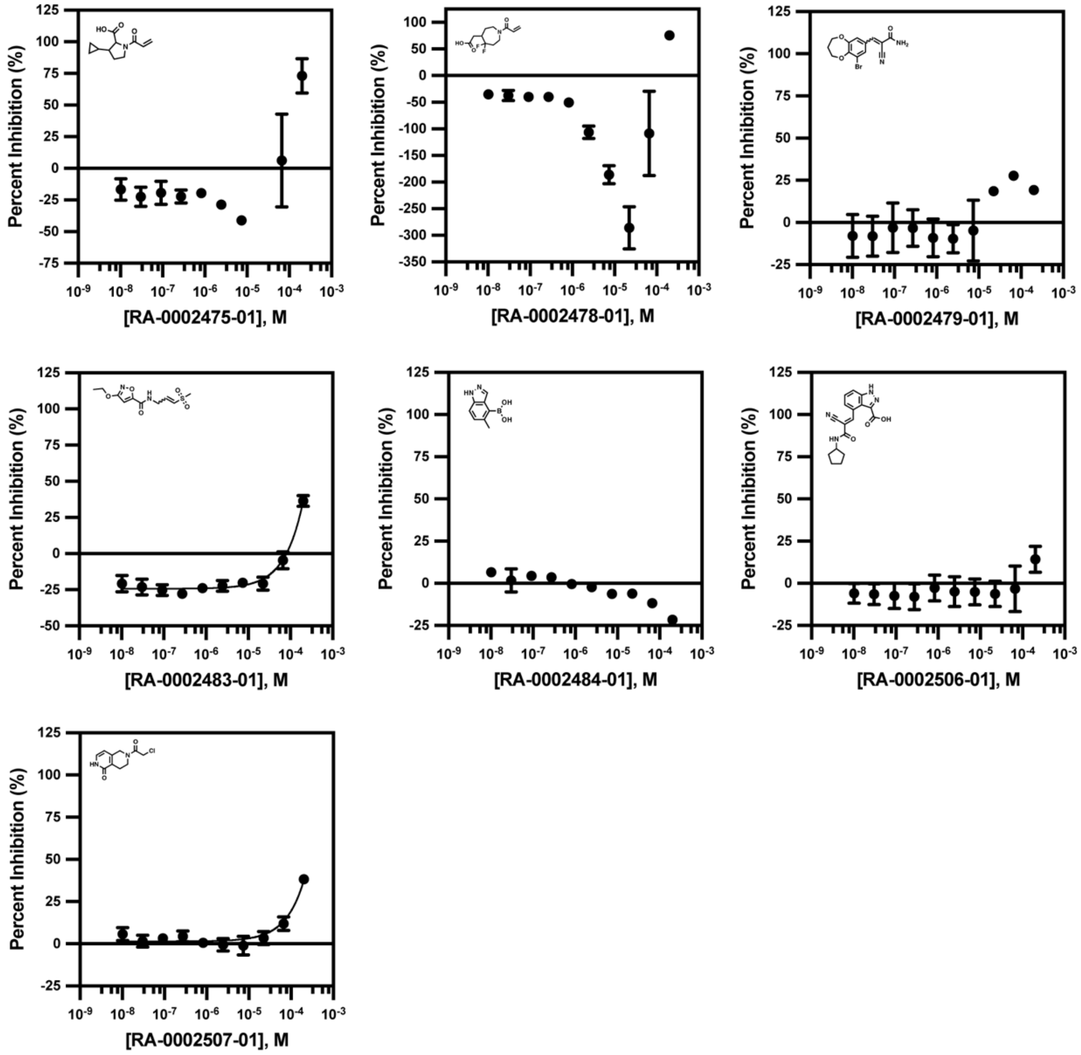


**Figure S5. Dose-response validation of 43 repurchased active compounds.** 43 repurchased compounds were tested in dose response format: 3-fold dilutions and 10 dilutions, beginning from 200 μM (n=3).

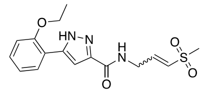

**RA-0002034**

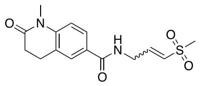

**RA-0002352**

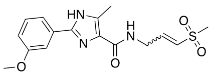

**RA-0002482**

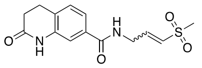

**RA-0002481**

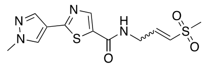

**RA-0002056**

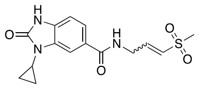

**RA-0002505**

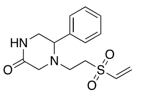

**RA-0002048**

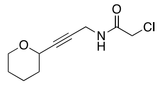

**RA-0002043**

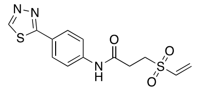

**RA-0002054**

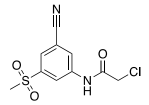

**RA-0002039**

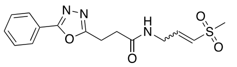

**RA-0002057**

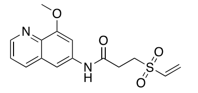

**RA-0002051**

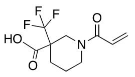

**RA-0002058**

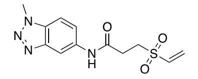

**RA-0002055**

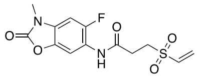

**RA-0002038**

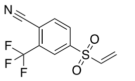

**RA-0002037**

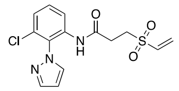

**RA-0002052**

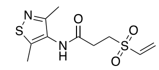

**RA-0002047**

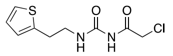

**RA-0002044**

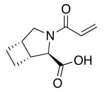

**RA-0002035**

**Figure S6.** Chemical structures of 20 compounds with $IC_{50}$s < 20 μM.

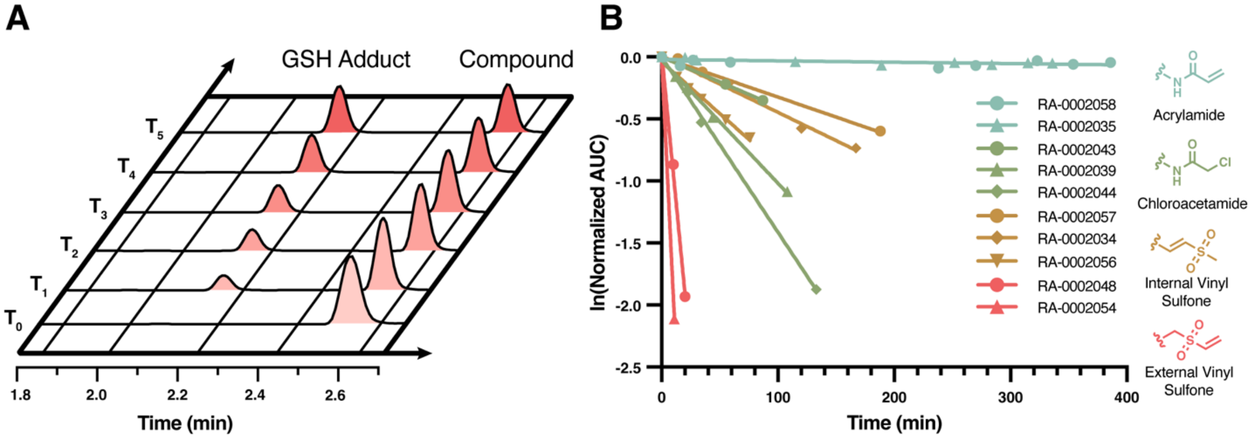


**Figure S7. GSH reactivity of active compounds.** (A) Principle of GSH assay; reaction of RA-0002056 with GSH over time. (B) Natural logs of normalized AUCs to rhodamine B internal standard plotted over time. A linear regression was used to calculate a GSH $t_{1/2}$. Compounds are colored by warhead type; acrylamides (blue), chloroacetamides (green), internal vinyl sulfones (yellow), and external vinyl sulfones (red).

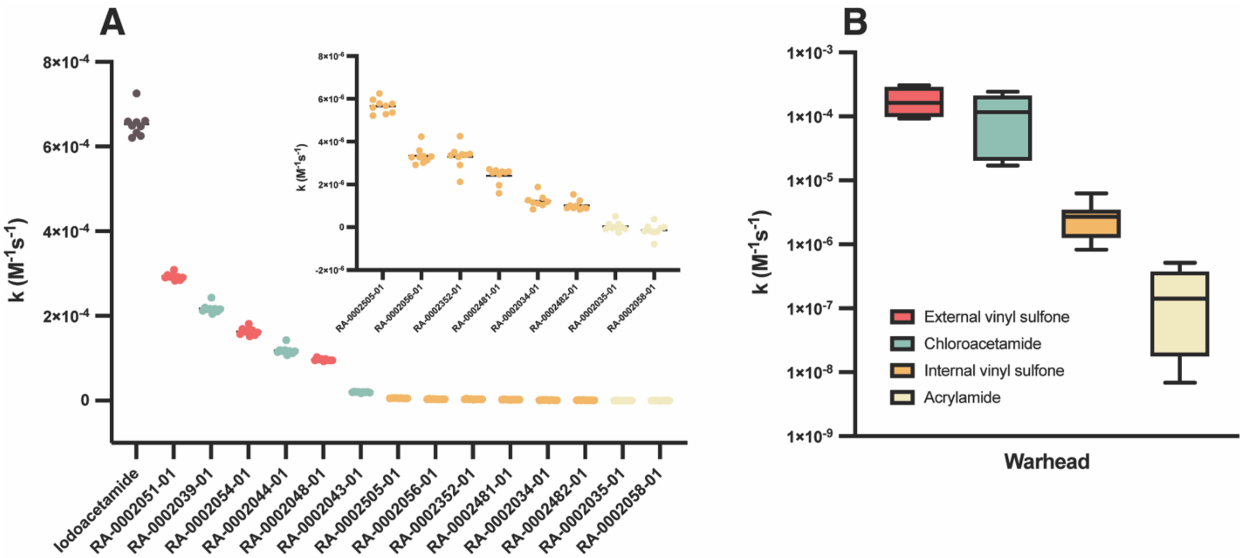


**Figure S8. DTNB thiol reactivity.** (A) Calculated reactivity rate for covalent fragments tested (n=9). Compounds are colored by warhead type; iodoacetamide (grey), acrylamides (yellow), chloroacetamides (blue), internal vinyl sulfones (orange), and external vinyl sulfones (red). (B) Average and standard deviations in DTNB assay for each warhead type tested.

**Table S3.** Thiol reactivity of covalent fragments.

| Compound | RA ID | UNC ID | Warhead | $IC_{50}$ (µM) | GSH $t_{1/2}$ (min) | k (M-1s-1) | Std Dev. | $R^2$ |
|---|---|---|---|---|---|---|---|---|
| 75 | RA-0002034 | UNC10307356 | Internal vinyl sulfone | 0.20 ± 0.06 | 130 | 1.10E-06 | 4.93E-07 | 0.957 |
| 18 | RA-0002352 | UNC10309756 | Internal vinyl sulfone | 0.39 ± 0.08 | | 3.30E-06 | 5.64E-07 | 0.99 |
| 10 | RA-0002482 | UNC10307390 | Internal vinyl sulfone | 0.59 ± 0.09 | | 1.03E-06 | 2.28E-07 | 0.857 |
| 16 | RA-0002481 | UNC10307388 | Internal vinyl sulfone | 1.0 ± 0.2 | | 2.41E-06 | 3.77E-07 | 0.986 |
| 21 | RA-0002056 | UNC10307194 | Internal vinyl sulfone | 1.2 ± 0.5 | 84 | 5.65E-06 | 3.33E-07 | 0.995 |
| 15 | RA-0002505 | UNC10307358 | Internal vinyl sulfone | 1.2 ± 0.5 | | 3.34E-06 | 3.86E-07 | 0.997 |
| 11 | RA-0002048 | UNC10307416 | External vinyl sulfone | 1.6 ± 0.2 | 7.2 | 9.68E-05 | 2.97E-06 | 0.999 |
| 46 | RA-0002043 | UNC10308193 | Chloroacetamide | 1.7 ± 0.1 | 260 | 1.96E-05 | 1.20E-06 | 0.993 |
| 29 | RA-0002054 | UNC10307220 | External vinyl sulfone | 3.4 ± 0.2 | 3.7 | 1.63E-04 | 8.96E-06 | 0.999 |
| 4 | RA-0002039 | UNC10308776 | Chloroacetamide | 4.5 ± 2.3 | 72 | 2.17E-04 | 1.05E-05 | 0.998 |
| 56 | RA-0002057 | UNC10307192 | Internal vinyl sulfone | 8.4 ± 1.5 | 200 | | | |
| 30 | RA-0002051 | UNC10307318 | External vinyl sulfone | 8.4 ± 0.3 | | 2.93E-04 | 7.63E-06 | 0.998 |
| 70 | RA-0002058 | UNC10305373 | Acrylamide | 8.6 ± 2.8 | >1400 | -9.72E-08 | 1.02E-07 | 0.087 |
| 66 | RA-0002055 | UNC10307210 | External vinyl sulfone | 9.1 ± 0.6 | | | | |
| 51 | RA-0002038 | UNC10309737 | External vinyl sulfone | 11 ± 2 | | | | |
| 107 | RA-0002037 | UNC10309765 | External vinyl sulfone | 12 ± 2 | | | | |
| 124 | RA-0002052 | UNC10307276 | External vinyl sulfone | 16 ± 2 | | | | |
| 99 | RA-0002047 | UNC10307486 | External vinyl sulfone | 17 ± 1 | | | | |
| 83 | RA-0002044 | UNC10307885 | Chloroacetamide | 18 ± 1 | 33 | 1.18E-04 | 1.01E-05 | 0.998 |
| 52 | RA-0002035 | UNC10305573 | Acrylamide | 20 ± 7 | >1400 | 4.40E-08 | 2.13E-07 | 0.291 |

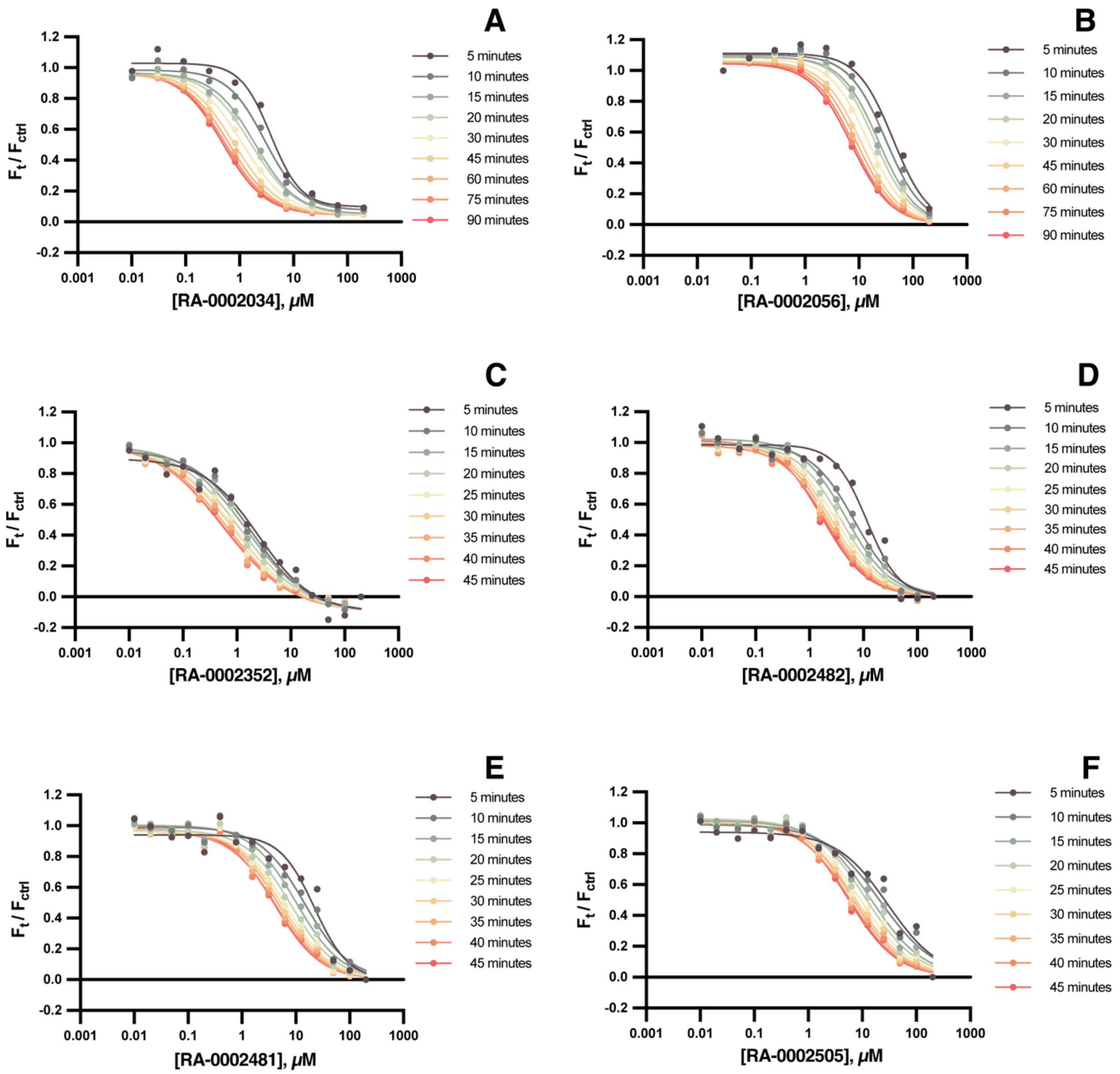


**Figure S9. Time-dependent $IC_{50}$ experiments.** Active compounds (A) RA-0002034, (B) RA-0002056, (C) RA-0002352, (D) RA-0002482, (E) RA-0002481, and (F) RA-0002505 were tested in dose-response format, calculating $IC_{50}$s at increasing timepoints.

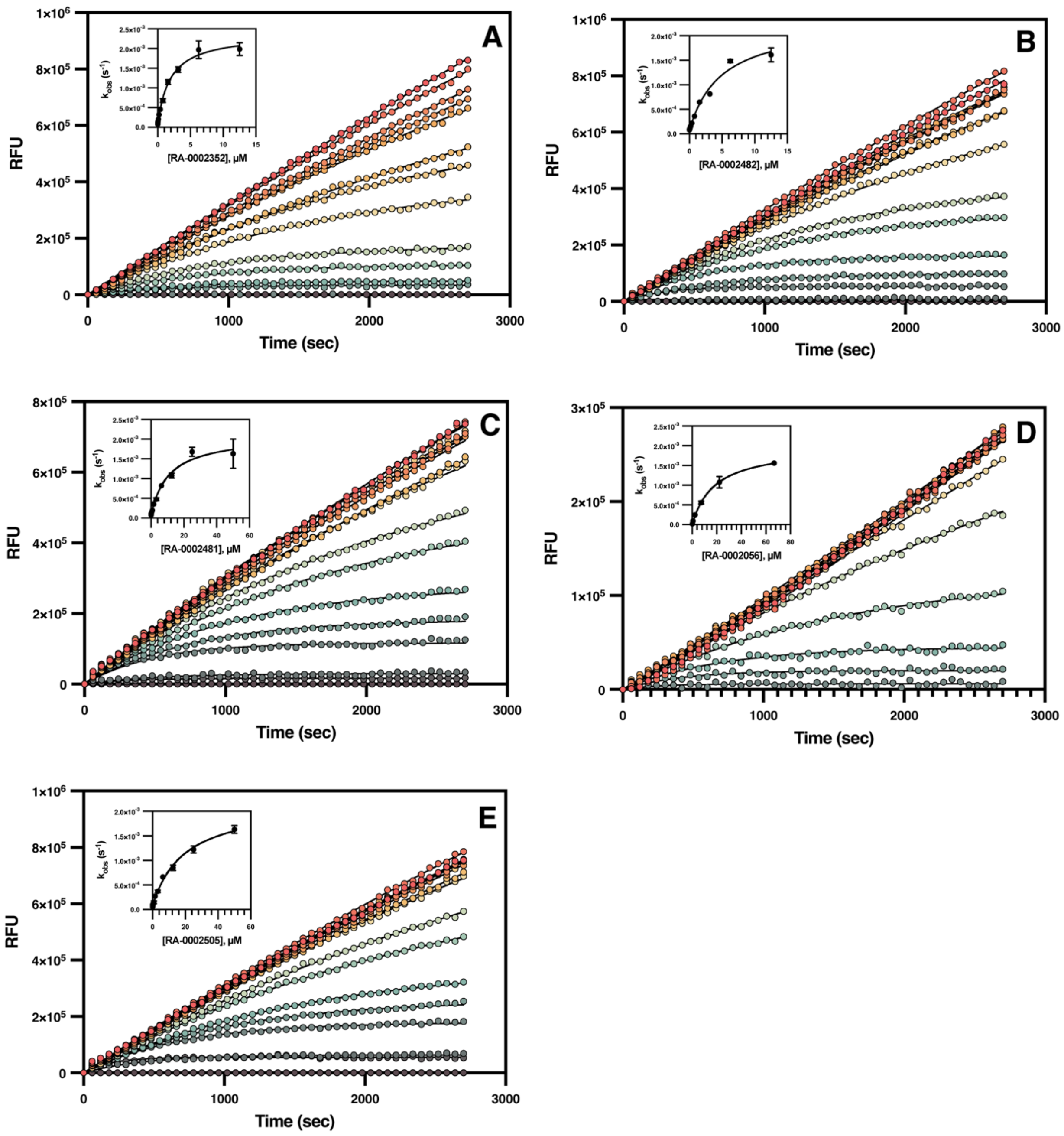


**Figure S10. Time-dependent inhibition for compounds 18, 10, 16, 21, and 15.** CHIKV nsP2pro is inhibited in a time-dependent manner. 2-fold serial dilutions of (A) RA-0002352, (B) RA-0002482, (C) RA-0002481, (D) RA-0002056, and (E) RA-0002505 were used to determine $K_I$, $k_{inact}$, and second order rate constant $k_{inact}/K_I$ (n=3).

**Scheme S1.** Synthesis of CHIKV nsP2pro covalent hit RA-0002034 **(75)**

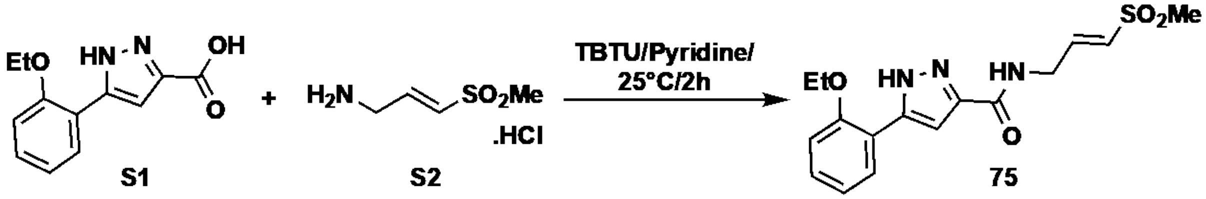


**RA-0002034 (75) Synthetic Procedure.** The CHIKV nsP2pro covalent hit RA-0002034 (75) was obtained by performing an acid-amine coupling reaction between 5-(2-ethoxyphenyl)-1H-pyrazole-3-carboxylic acid (S1) and (E)-3-(methylsulfonyl) prop-2-en-1-amine hydrochloride (S2). Compounds S1 and S2 were synthesized by performing 4-step and 3-step reactions respectively (discussed under Schemes S2 and S3). Discussed in the supporting information are the synthetic protocols and analytical data of the intermediates and final compounds for acid S1, and amine S2. To synthesize RA-0002034 (75), amine S2 (89 mg, 0.52 mmol, 1.2 eq.) was added to a stirred solution of S1 (100 mg, 0.43 mmol, 1.0 eq.) and TBTU (207 mg, 0.65 mmol, 1.5 eq.) in pyridine (3 mL), and the reaction was stirred at 25 ºC for 2h. On completion of the reaction (based on TLC and LCMS), the reaction was poured into water and extracted with ethyl acetate. The combined organic layers were dried over anhydrous $Na_2SO_4$, filtered, and concentrated in vacuo to give the crude compound. Column chromatography (eluting with 0-100% EtOAc in hexanes), followed by preparative HPLC purification afforded (E)-5-(2-ethoxyphenyl)-N-(3-(methylsulfonyl)allyl)-1H-pyrazole-3-carboxamide, trifluoroacetate salt (75) as a white solid (140 mg, 70%). $^1$H NMR (DMSO-d6, 500 MHz): δ 8.62 (t, J = 5.9 Hz, 1H), 7.74 (dd, J = 7.5, 1.7 Hz, 1H), 7.35 (ddd, J = 8.7, 7.4, 1.7 Hz, 1H), 7.16 – 7.10 (m, 2H), 7.03 (t, J = 7.5 Hz, 1H), 6.82 (dt, J = 15.3, 4.4 Hz, 1H), 6.69 (dt, J = 15.3, 1.8 Hz, 1H), 4.17 (q, J = 7.0 Hz, 2H), 4.11 (ddd, J = 6.2, 4.2, 1.8 Hz, 2H), 3.01 (s, 3H), 1.41 (t, J = 6.9 Hz, 3H); $^{13}$C NMR (DMSO-d6, 126 MHz): δ 161.4, 155.0, 143.5, 130.0, 129.7, 127.8, 120.6, 112.8, 105.6, 63.7, 42.1, 40.0, 39.9, 39.7, 39.5, 39.4, 39.2, 39.0, 38.7, 14.6. HRMS (ESI) m/z: $[M+H]^+$ calculated for $C_{16}H_{20}N_3O_4S$: 350.1175; found 350.1172. HPLC purity >99%.

**Scheme S2.** Synthesis of 5-(2-ethoxyphenyl)-1*H*-pyrazole-3-carboxylic acid **(S1)**

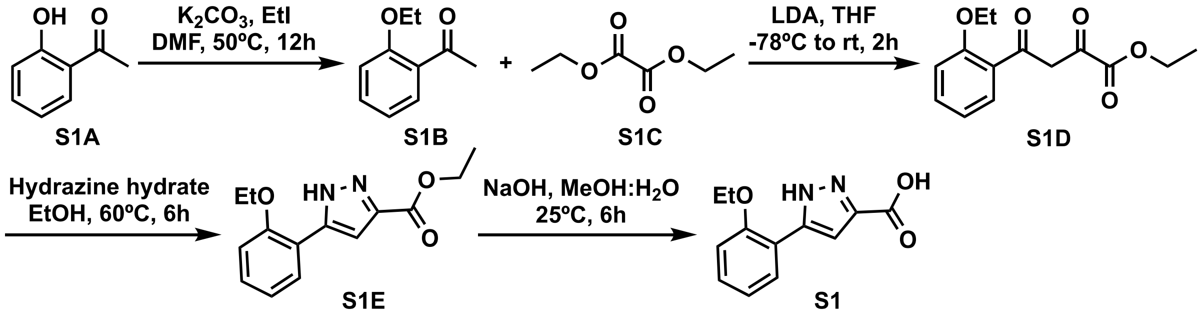


**1-(2-ethoxyphenyl) ethan-1-one (S1B)**

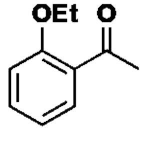


To a stirred solution of 1-(2-hydroxyphenyl)ethan-1-one **S1A** (5.0 g, 36.7 mmol, 1.0 eq.) in DMF (50 mL) at 25 °C was added $K_2CO_3$ (7.6 g, 44.1 mmol, 1.2 eq.) and then ethyl iodide (6 mL, 73.5 mmol, 2.0 eq.). Reaction mixture was stirred at 50 °C for 12h. After completion of the reaction (based on TLC and LCMS), the reaction mixture was poured into ice cold water. Organic layer was extracted with ethyl acetate. Combined organic layers were dried over anhydrous $Na_2SO_4$ and concentrated under vacuum. Column chromatography (eluting with 0-30% EtOAc in hexanes) afforded 1-(2-ethoxyphenyl)ethan-1-one **(S1B)** as a white solid (4 g, 66%). m/z $[M+H]^+$ 165.3

**Ethyl 4-(2-ethoxyphenyl)-2,4-dioxobutanoate (S1D)**

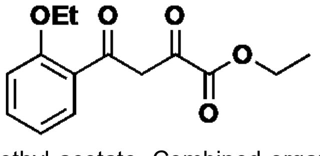


To a stirred solution of **S1B** (4 g, 24.4 mmol, 1.0 eq.) in THF (40 mL) was added 2M LDA solution (60 mL, 122 mmol, 1.0 eq.) dropwise at -78 °C and stirred at same temperature for 30 min. Then at the same temperature, diethyl oxalate (**S1C**) (5.3 g, 36.6 mmol, 1.5 eq.) was added and the reaction mixture was stirred at 25 °C for 2h. On completion of the reaction (based on TLC and LCMS analyses), the reaction mixture was acidified with 1N HCl until pH ~ 3 and the organic layer was extracted with ethyl acetate. Combined organic layers were washed with brine, dried over anhydrous $Na_2SO_4$, and concentrated *in*

*vacuo*. Column chromatography (eluting with 2-5% EtOAc in hexanes) afforded ethyl 4-(2-ethoxyphenyl)-2,4-dioxobutanoate **(S1D)** as yellow solid (1.8 g, 28%).

**Ethyl 5-(2-ethoxyphenyl)-1H-pyrazole-3-carboxylate (S1E)**

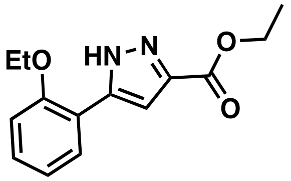


To a stirred solution of **S1D** (1.8 g, 6.8 mmol, 1.0 eq.) in EtOH (20 mL) was added hydrazine hydrate (50% by wt.) (0.4 mL, 8.2 mmol, 1.2 eq.) dropwise and the reaction was stirred at 60 °C for 6h. After completion of the reaction (confirmed by TLC and LCMS), the reaction mixture was concentrated under vacuum. Column chromatography (eluting with 15-25% EtOAc in hexanes) afforded ethyl 5-(2-ethoxyphenyl)-1H-pyrazole-3-carboxylate **(S1E)** as pale yellow solid (1 g, 56%). m/z $[M+H]^+$ 261.0.

**5-(2-ethoxyphenyl)-1H-pyrazole-3-carboxylic acid (S1)**

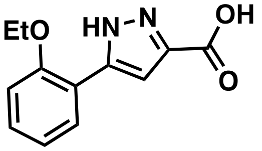


To a stirred solution of **S1E** (1 g, 3.8 mmol, 1.0 eq.) in MeOH (6 mL) and water (4 mL) was added NaOH (0.4 g, 7.7 mmol, 2.0 eq). The reaction was stirred at 25 °C for 6h. On completion of the reaction (based on TLC and LCMS analyses), the reaction mixture was concentrated under reduce pressure, residue acidified with 1N HCl until pH ~ 3. Organic layer was extracted with ethyl acetate, combined organic layers dried over anhydrous $Na_2SO_4$, and concentrated under reduce pressure to afford 5-(2-ethoxyphenyl)-1H-pyrazole-3-carboxylic acid **(S1)** as a light yellow solid (0.65 g, 73%). **$^1$H NMR** (DMSO-$d_6$, 400 MHz): $\delta$ 13.24 (s, 2H), 7.81 (dd, $J$ = 7.7, 1.7 Hz, 1H), 7.33 (ddd, $J$ = 8.3, 7.3, 1.7 Hz, 1H), 7.18 – 7.08 (m, 2H), 7.01 (td, $J$ = 7.5, 1.1 Hz, 1H), 4.15 (q, $J$ = 6.9 Hz, 2H), 1.40 (t, $J$ = 6.9 Hz, 3H); **$^{13}$C NMR** (DMSO-$d_6$, 100 MHz): $\delta$ 155.1, 129.5, 127.6, 120.5, 112.7, 108.2, 63.6, 14.6. m/z $[M+H]^+$ 233

**Scheme S3.** Synthesis of (*E*)-3-(methylsulfonyl)prop-2-en-1-amine hydrochloride **(S2)**

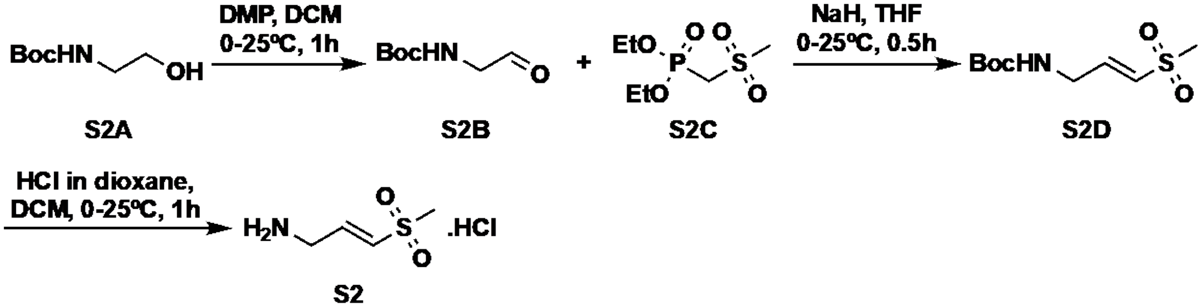


***tert*-Butyl (2-oxoethyl)carbamate (S2B)**

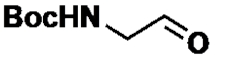


To a stirred solution of tert-butyl (2-hydroxyethyl) carbamate **S2A** (5.0 g, 31.05 mmol, 1.0 eq.) in DCM (50 mL), Dess-martin periodinane (19.75 g, 46.58 mmol, 1.5 eq.) was added at 0 °C. The reaction was stirred at 25 °C for 1h. On completion of the reaction (based on TLC and LCMS), the reaction mixture was diluted with $NaHCO_3$ solution (100 mL) and extracted with DCM (2 x 100 mL). The organic layers were combined and dried over anhydrous $Na_2SO_4$, filtered, and concentrated under reduced pressure to afford *tert*-butyl (2-oxoethyl)carbamate (**S2B**) (4.0 g) as a colorless liquid. The crude was directly used for next step without further purification.

***tert*-butyl (*E*)-(3-(methylsulfonyl) allyl) carbamate (S2D)**

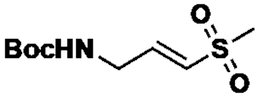


To a stirred solution of diethyl ((methylsulfonyl)methyl)phosphonate **S2C** (2.5 g, 11.30 mmol, 0.45 eq.) in THF (40 mL), NaH (60% dispersion in mineral oil) (1.5 g, 37.73 mmol, 1.5 eq.) was added at 0 °C in portions. The reaction mixture was stirred at 0 °C for 15 min. Then *tert*-butyl (2-oxoethyl)carbamate (**S2B**) (4.0 g, 25.15 mmol, 1.0 eq.) was added at 0 °C and the reaction was stirred at 25 °C for 15 min. On completion of the reaction (observed by TLC), the reaction mixture was poured into water (100 mL) and extracted with ethyl acetate (2 x 100 mL). The combined organic layers were dried over anhydrous $Na_2SO_4$, filtered, and concentrated *in vacuo* to give the crude compound. Column chromatography (eluting with 0-30% EtOAc in hexanes) afforded *tert*-butyl (*E*)-(3-(methylsulfonyl) allyl) carbamate **(S2D)** as a pale yellow liquid (1.0 g, 17%).

**(*E*)-3-(methylsulfonyl) prop-2-en-1-amine hydrochloride (S2)**

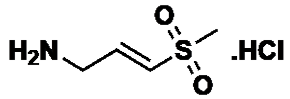


To a stirred solution of *tert*-butyl (*E*)-(3-(methylsulfonyl) allyl) carbamate (**S2D**) (1.0 g, 4.2 mmol, 1.0 eq.) in DCM (10 mL), 4M HCl in dioxane (10 mL) was added at 0 °C. The reaction mixture was stirred at 25 °C for 1h. On completion of the reaction (based on TLC), the reaction mixture was concentrated to give the crude compound. The resulting crude was purified by trituration using diethyl ether (20 mL), then dried under reduced pressure to afford (*E*)-3-(methylsulfonyl) prop-2-en-1-amine hydrochloride **(S2)** as a yellow solid (0.675 g, 93%). **$^1$H NMR** (DMSO-*d*6, 400 MHz): *δ* 8.51 (s, 2H), 7.02 (dt, *J* = 15.4, 1.7 Hz, 1H), 6.76 (dt, *J* = 15.5, 5.6 Hz, 1H), 3.72 (d, *J* = 5.7 Hz, 2H), 3.04 (s, 3H); **$^{13}$C NMR** (DMSO-*d*6, 100 MHz): *δ* 138.1, 133.4, 42.2, 38.4. m/z $[M+H]^+$ 136

**NMR Spectra**

**Figure S11A.** $^1$H NMR of compound **S1**

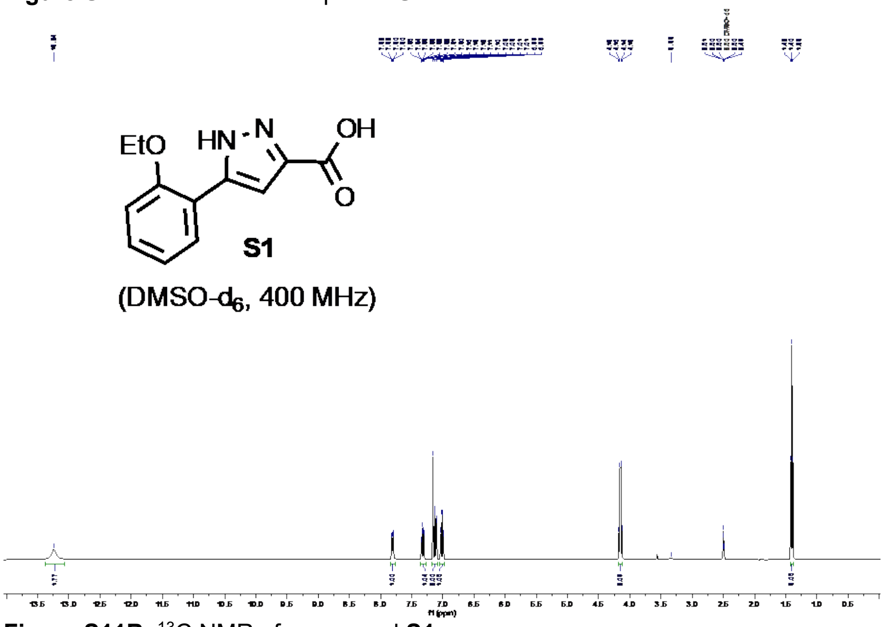


**Figure S11B.** $^{13}$C NMR of compound **S1**

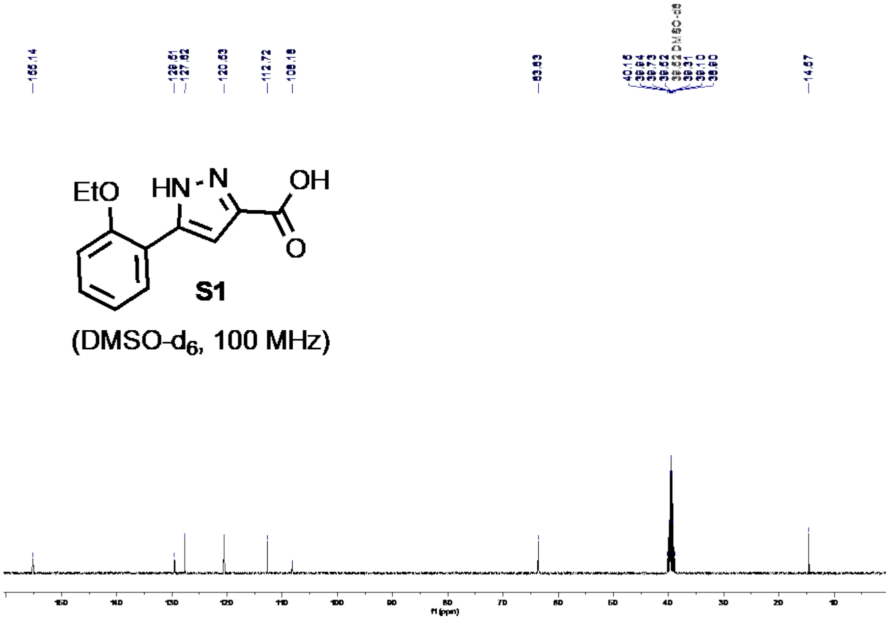

**Figure S11C.** $^{1}$H NMR of compound **S2**

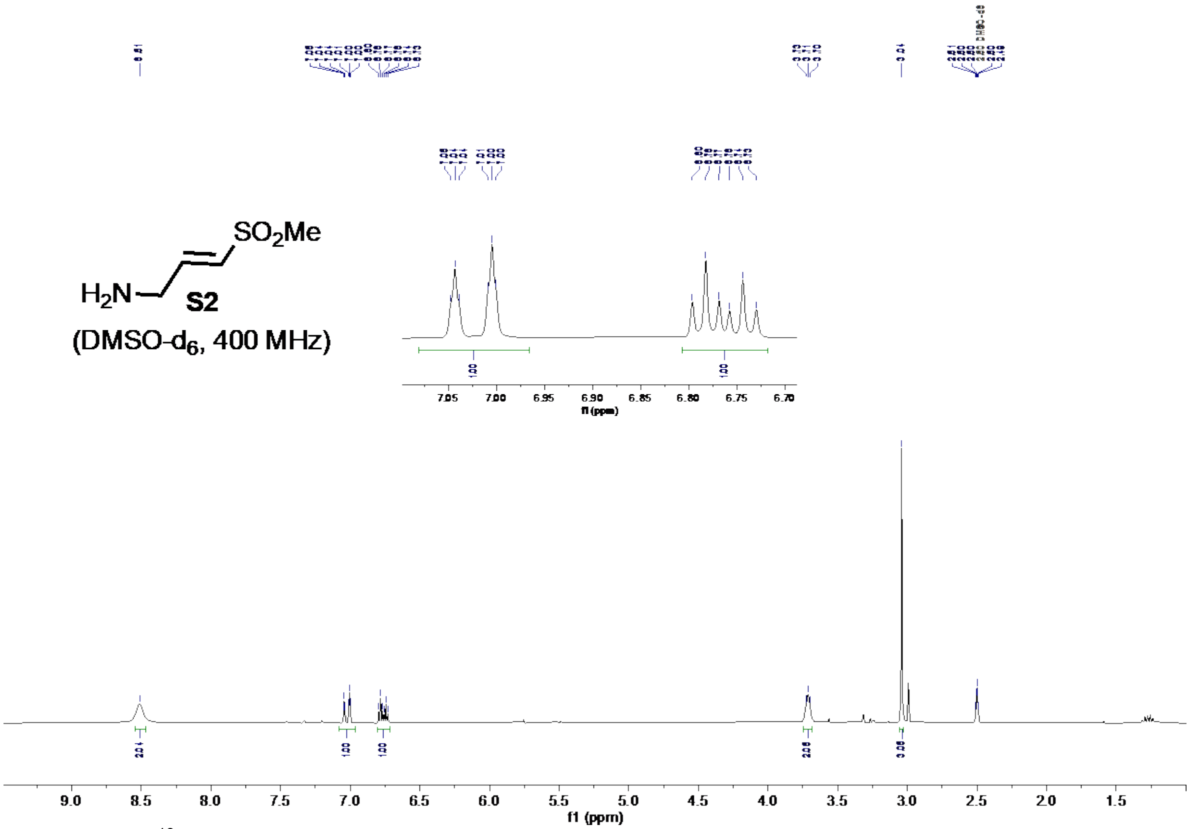


**Figure S11D.** $^{13}$C NMR of compound **S2**

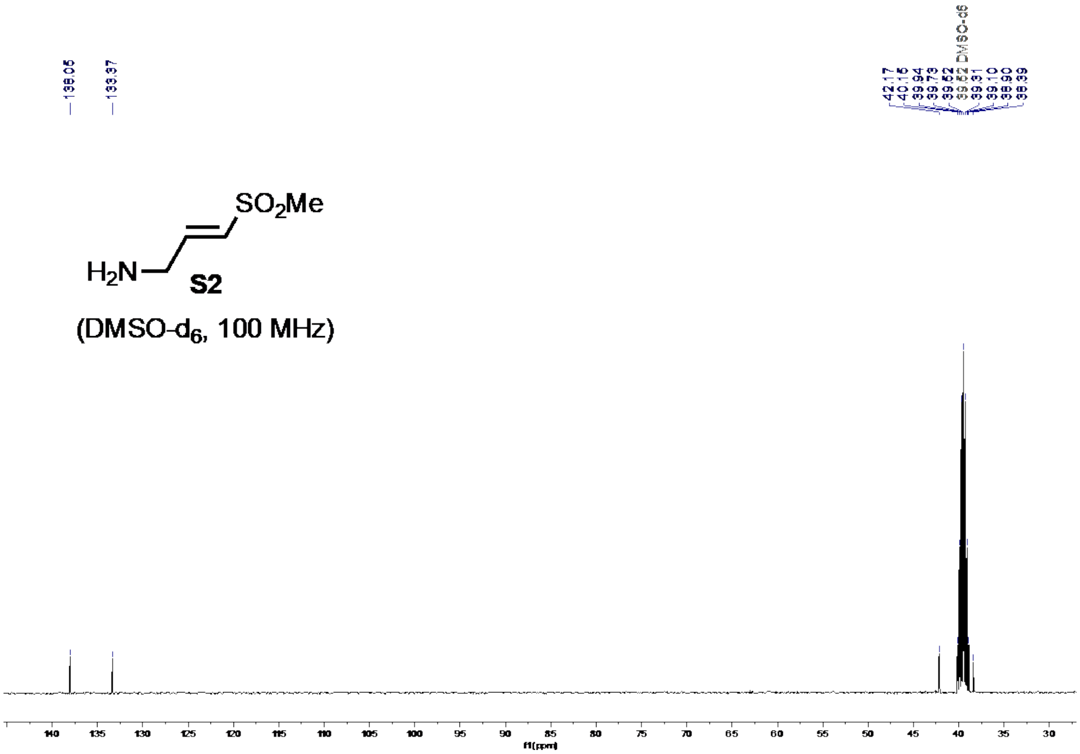

**Figure S11E.** $^{1}$H NMR of compound **75**

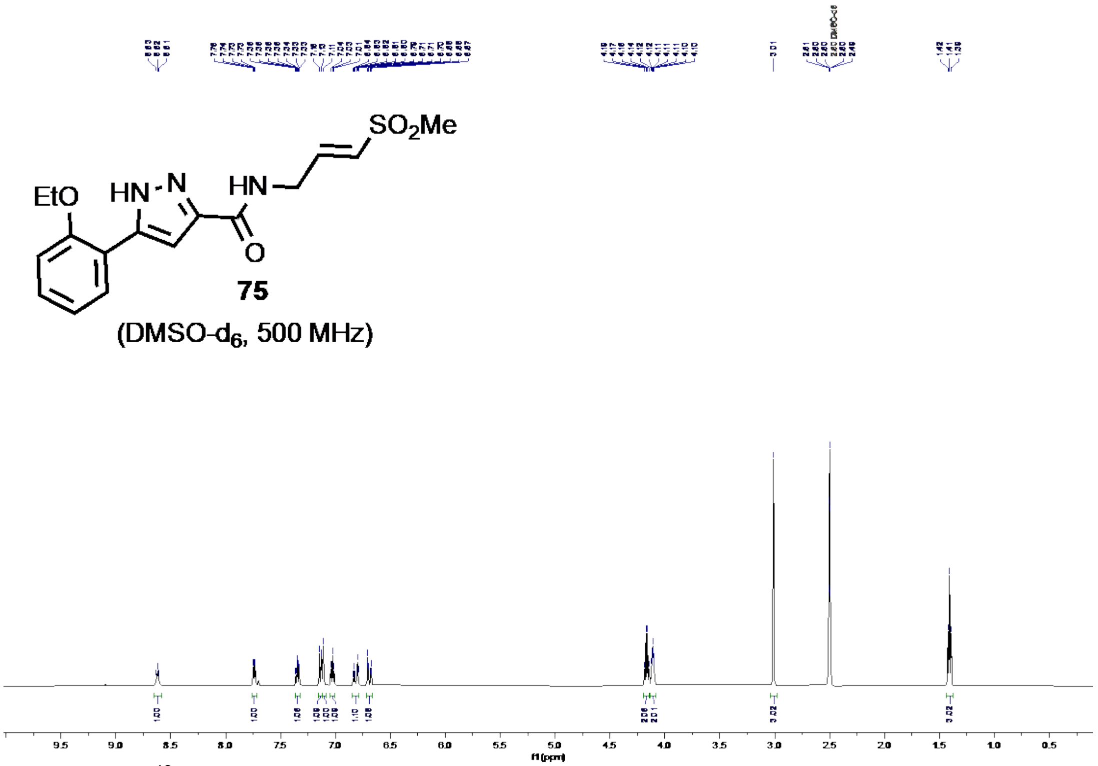


**Figure S11F.** $^{13}$C NMR of compound **75**

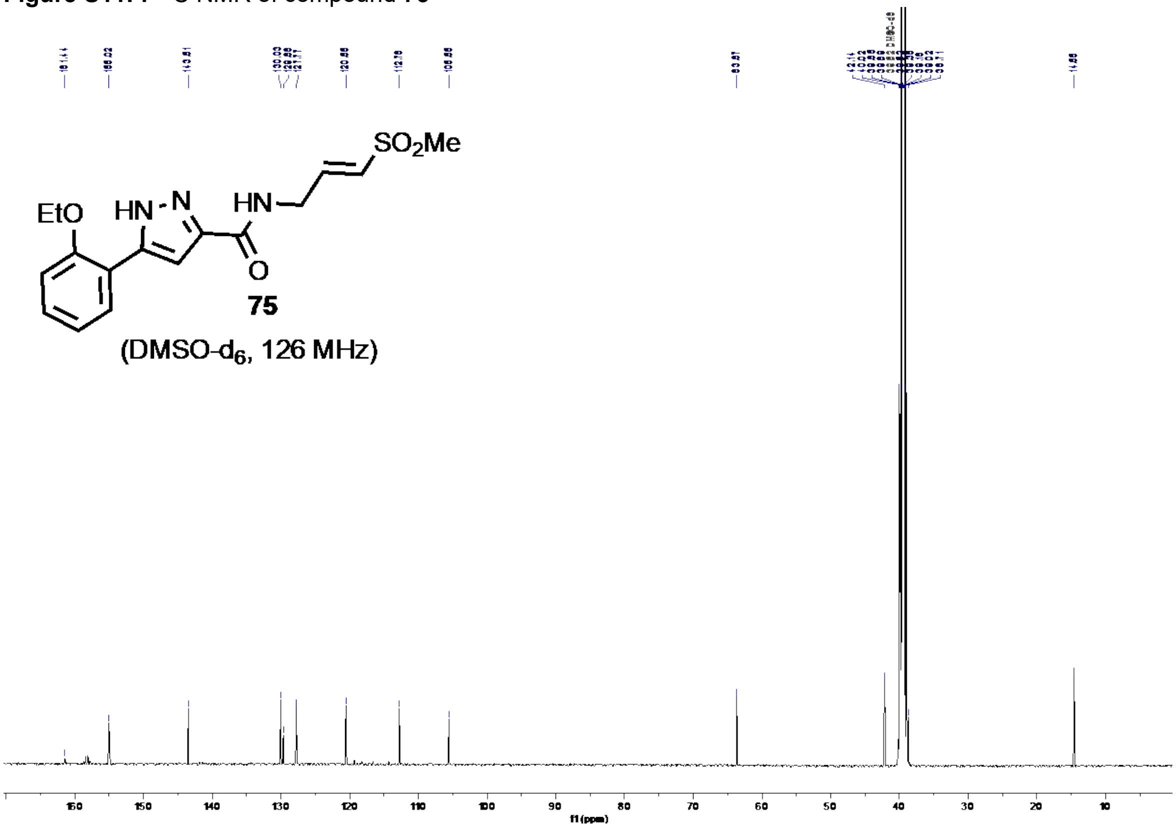

**Figure S11G.** LCMS and HPLC purity of compound **75 (RA-0002034)**

### LCMS Sample Purity - Detailed Report

Agilent

| | | | |
|---|---|---|---|
| **Result set name:** | SingleSample | | |
| **Data file:** | AG-02-045-P1-V43.dx | **Project name:** | SGC-UNC |
| **Sample name:** | AG-02-045-P1-V43 | **Operator:** | SYSTEM (SYSTEM) |
| **Instrument:** | LCMS G6160A | **Acquired on:** | 2023-08-03 12:23:53-04:00 |
| **Inj. volume:** | 4.000 µL | **Location:** | p2-a6 |
| **Acq. method:** | SGC_AH_opt1_4uL.amx | | |
| **Processing method:** | LC_MS Sample Purity_SGC.pmx | | |

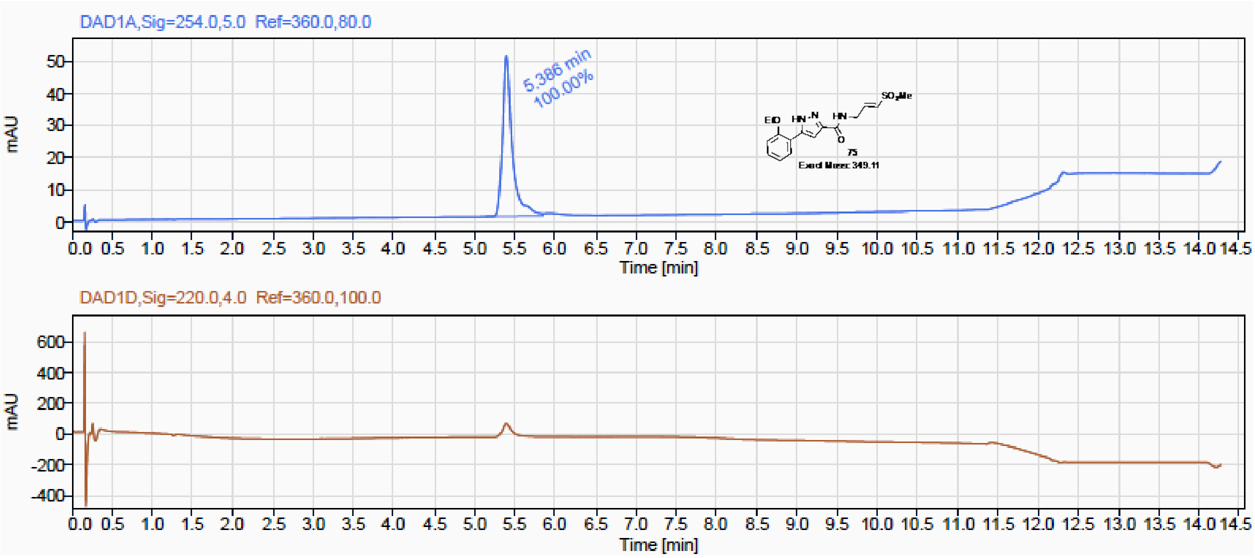


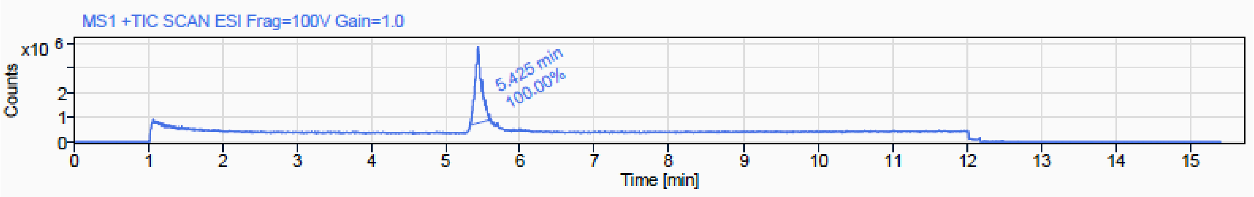

### Peak Results (Area Percent at least 1%)

Signal Name DAD1A

| RT (min) | Signal description | m/z | Purity UV(‰) / MS(%) | Area | Area% |
|---|---|---|---|---|---|
| 5.386 | DAD1A,Sig=254.0,5.0 Ref=360.0,80.0 | | | 421.6 | 100.00 |

Signal Name MS1 +TIC SCAN ESI Frag=100V Gain=1.0

| RT (min) | Signal description | m/z | Purity UV(‰) / MS(%) | Area | Area% |
|---|---|---|---|---|---|
| 5.425 | MS1 +TIC SCAN ESI Frag=100V Gain=1.0 | | | 19616009.4 | 100.00 |

### Peak Details (Area Percent at least 1%)

**Signal Description:** MS1 +TIC SCAN ESI Frag=100V Gain=1.0

**Peak RT:** 5.425 min **Area %:** 100.00%

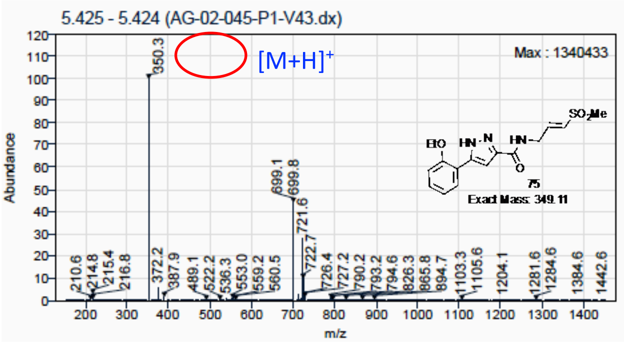


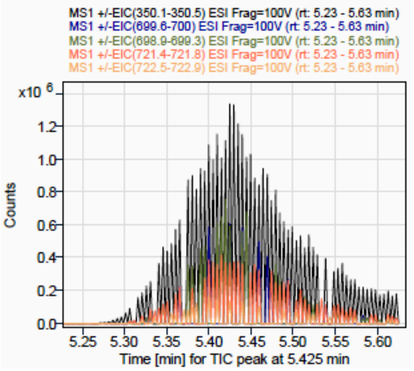


**Figure S11H.** HRMS spectrum of compound **75**

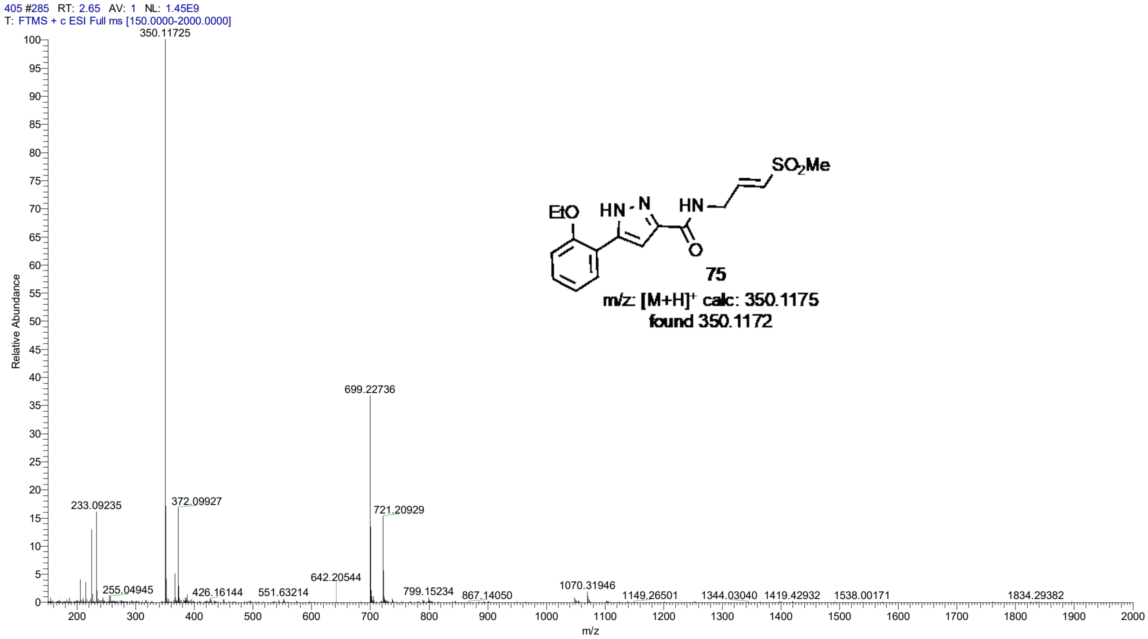

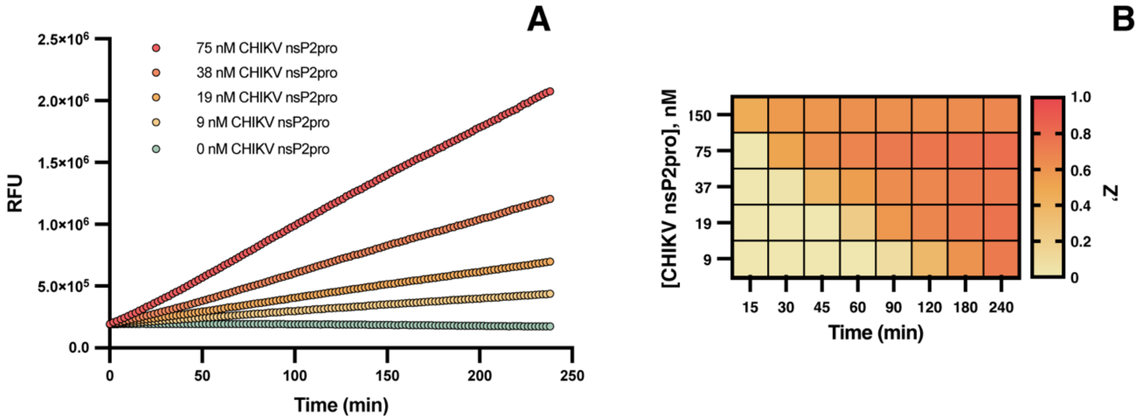


**Figure S12. CHIKV nsP2pro enzymatic assay re-optimization.** (A) CHIKV nsP2pro concentration was serially diluted (2-fold) beginning from 150 nM. (B) Z' scores were calculated over time using 0 nM CHIKV nsP2pro as the low signal control.

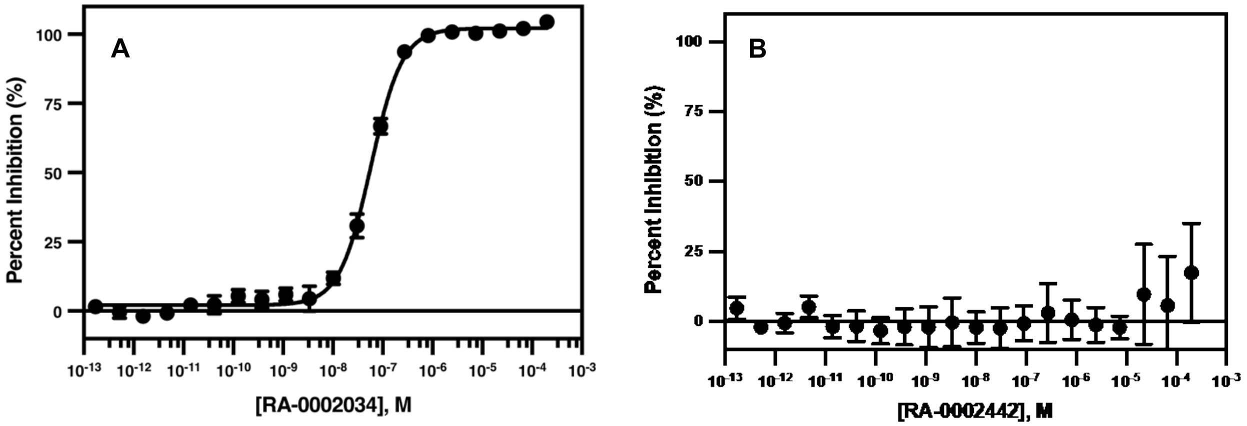


**Figure S13. Dose-response validation of resynthesized RA-0002034 and cyclic RA-0002442.** Resynthesized RA-0002034 (A) or cyclic analog RA-0002442 (B) were serially diluted (3-fold) over 20 dilutions beginning from 200 µM and tested in the enzymatic activity assay. The resulting $IC_{50}$ for RA-0002034 was 58 ± 17 nM, and >200 µM for RA-0002442 (n=3, with data points representing mean and standard deviation).

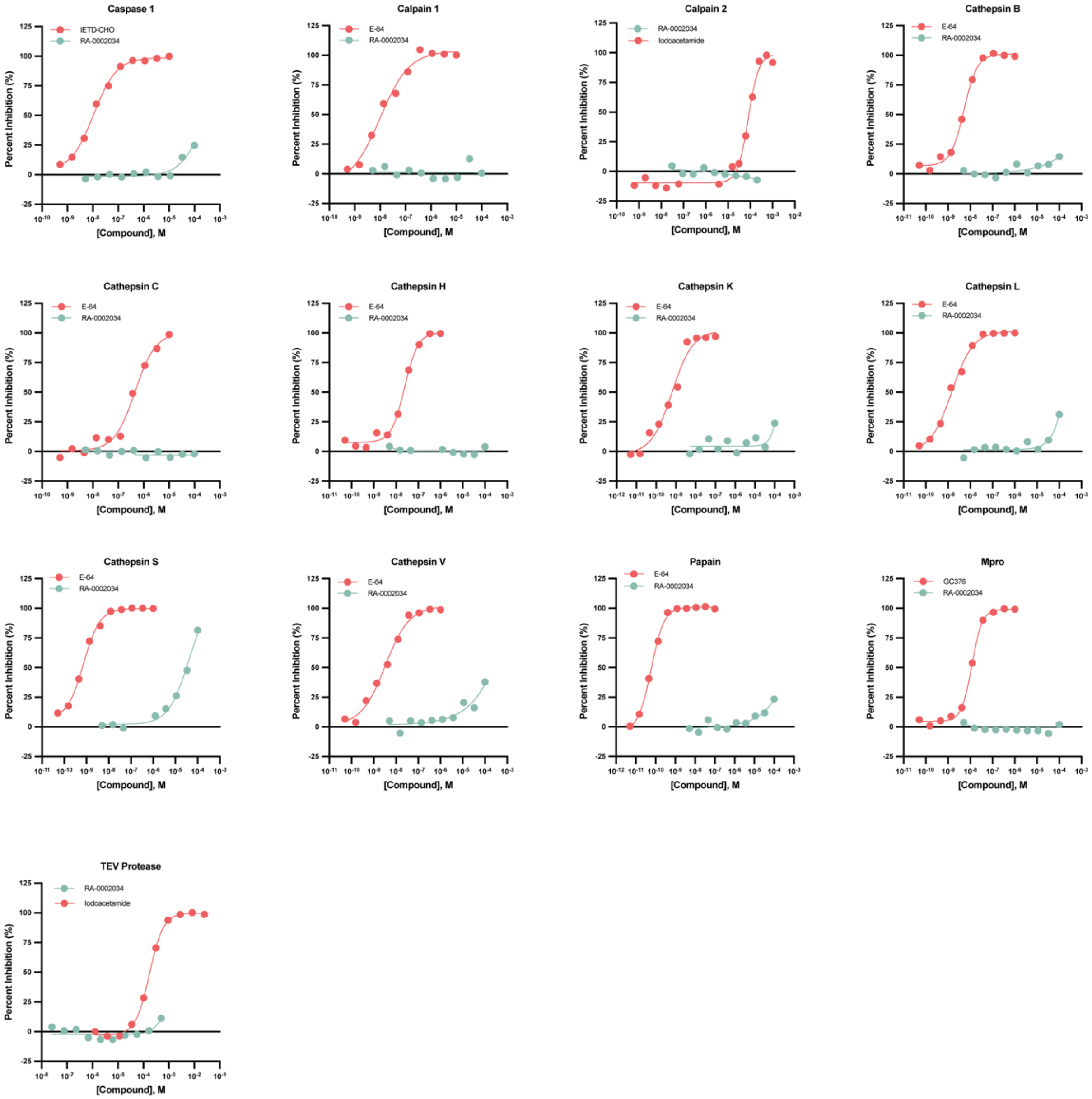


**Figure S14. Cysteine protease selectivity panel.** RA-0002034 was tested in dose-response format against 13 diverse cysteine proteases (n=1); caspase 1, calpain 1, calpain 2, cathepsin B, cathepsin C, cathepsin H, cathepsin K, cathepsin L, cathepsin S, cathepsin V, papain, SARS-CoV-2 Mpro, and TEV protease.

**Table S4.** Cysteine protease panel.

| Enzyme | Substrate | Substrate (μM) | Control | RA-0002034 $IC_{50}$ (μM) | Control $IC_{50}$ (μM) |
|---|---|---|---|---|---|
| Caspase 1 | Ac-LEHD-AMC | 5 | IETD-CHO | >100 | 0.011 |
| Calpain 1 | N-Succinyl-LY-AMC | 10 | E-64 | >100 | 0.010 |
| Calpain 2 | EDANS-PLFAERK-DABCYL | 30 | Iodoacetamide | >100 | 80 |
| Cathepsin B | Z-FR-AMC | 10 | E-64 | >100 | 0.0052 |
| Cathepsin C | H-GR-AMC | 10 | E-64 | >100 | 0.46 |
| Cathepsin H | H-Arg-AMC | 10 | E-64 | >100 | 0.025 |
| Cathepsin K | Z-FR-AMC | 5 | E-64 | >100 | 0.00064 |
| Cathepsin L | Z-FR-AMC | 10 | E-64 | >100 | 0.0015 |
| Cathepsin S | Z-FR-AMC | 10 | E-64 | 40 | 0.00073 |
| Cathepsin V | Z-FR-AMC | 10 | E-64 | >100 | 0.0033 |
| Papain | Z-FR-AMC | 10 | E-64 | >100 | 0.000060 |
| Mpro | NH2-C(EDANS)VNSTQSGLRK(DABCYL)M-COOH | 5 | GC376 | >100 | 0.012 |
| TEV protease | 5-FAM-ENLYFQG-QXL520 | 1x | Iodoacetamide | >200 | 170 |

AMC – 7-amino-4-methylcoumarin, EDANS – (5-((2-Aminoethyl) amino)naphthalene-1-sulfonic acid), DABCYL - N-[4-(4-dimethylamino)phenylazo]benzoic acid, FAM – Carboxyfluorescein.

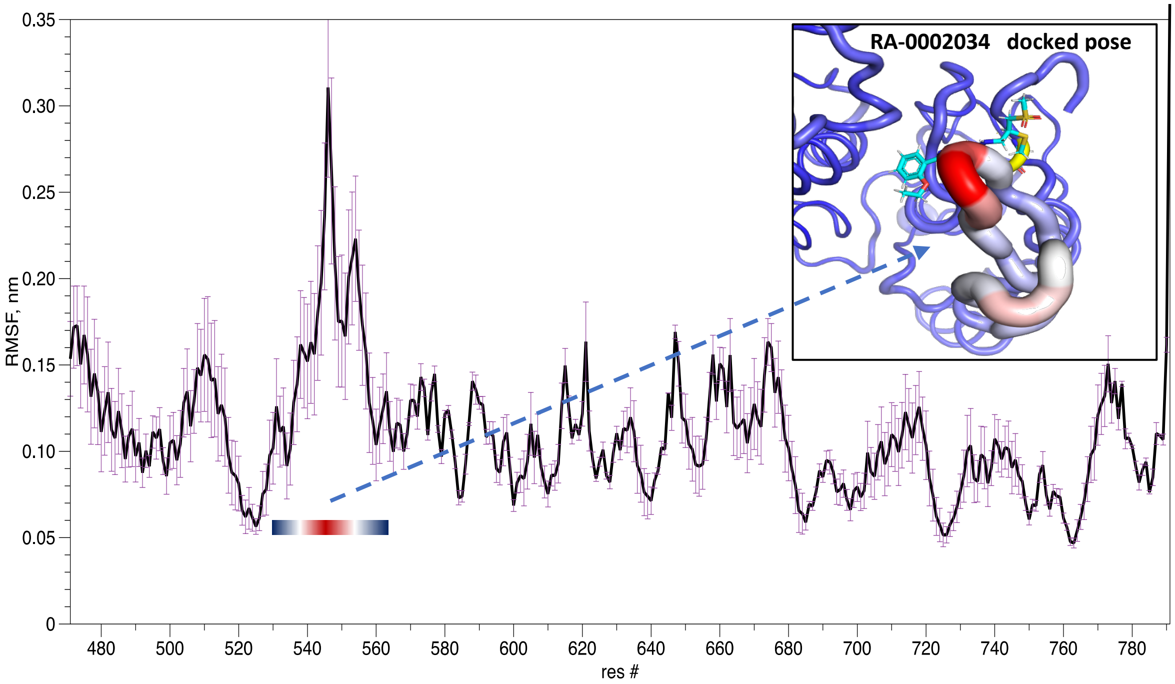


**Figure S15. CHIKV nsP2pro MD simulations.** nsP2 protease residue Real-mean-square-fluctuation (RMSF) values calculated from MD simulates. The inset panel illustrates the docking model of RA-0002034 nsP2 complex, where RMSF values are overlaid on the protein structure. Thicker tubes represent regions with higher backbone fluctuations, with red representing higher fluctuation and blue representing lower fluctuation.

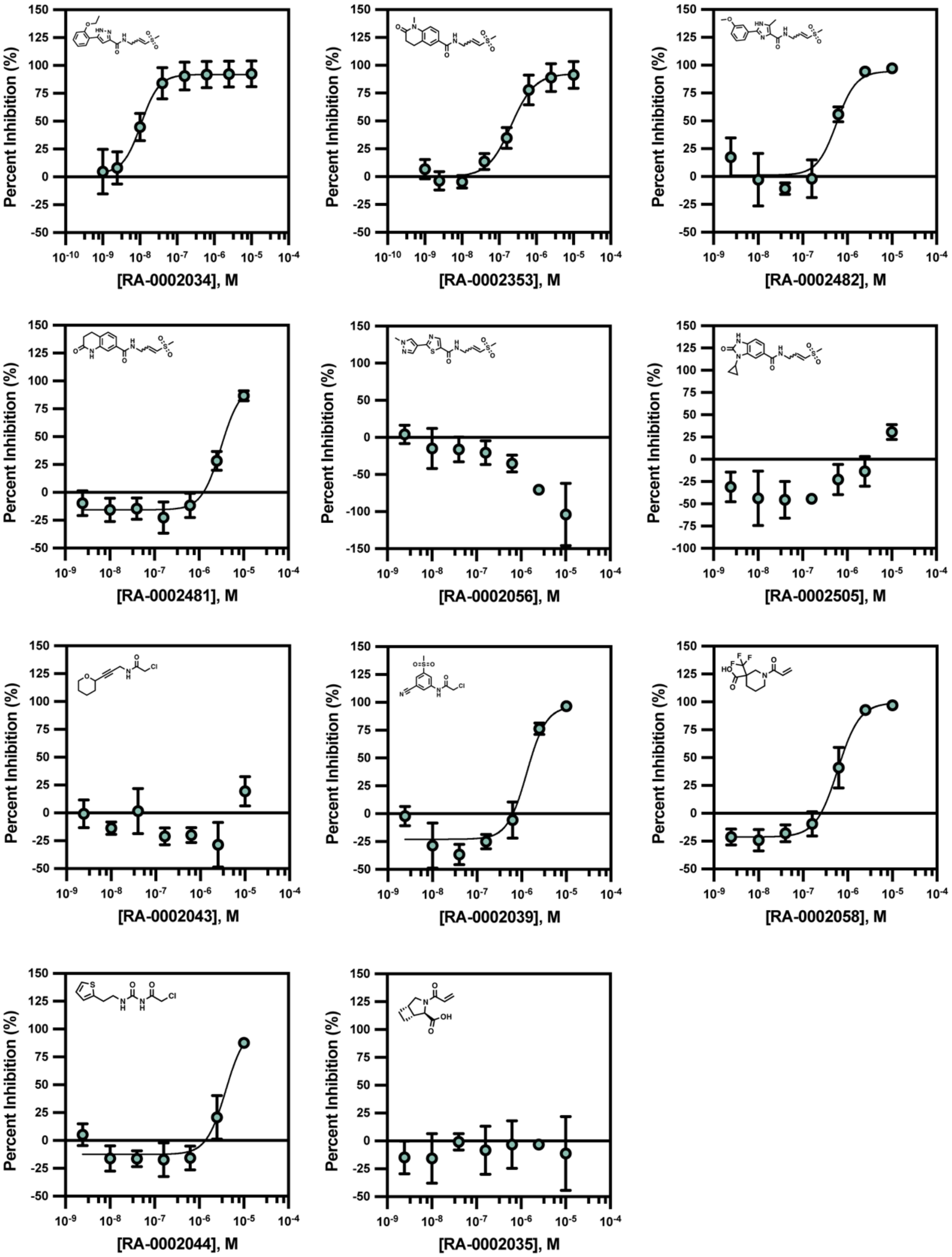


**Figure S16. Covalent fragment CHIKV nLuc dose-response results.** Active compounds in CHIKV nsP2pro activity assay were tested in a CHIKV nLuc reporter assay in dose-response format, using 4-fold dilutions beginning from 10 μM (n=3).

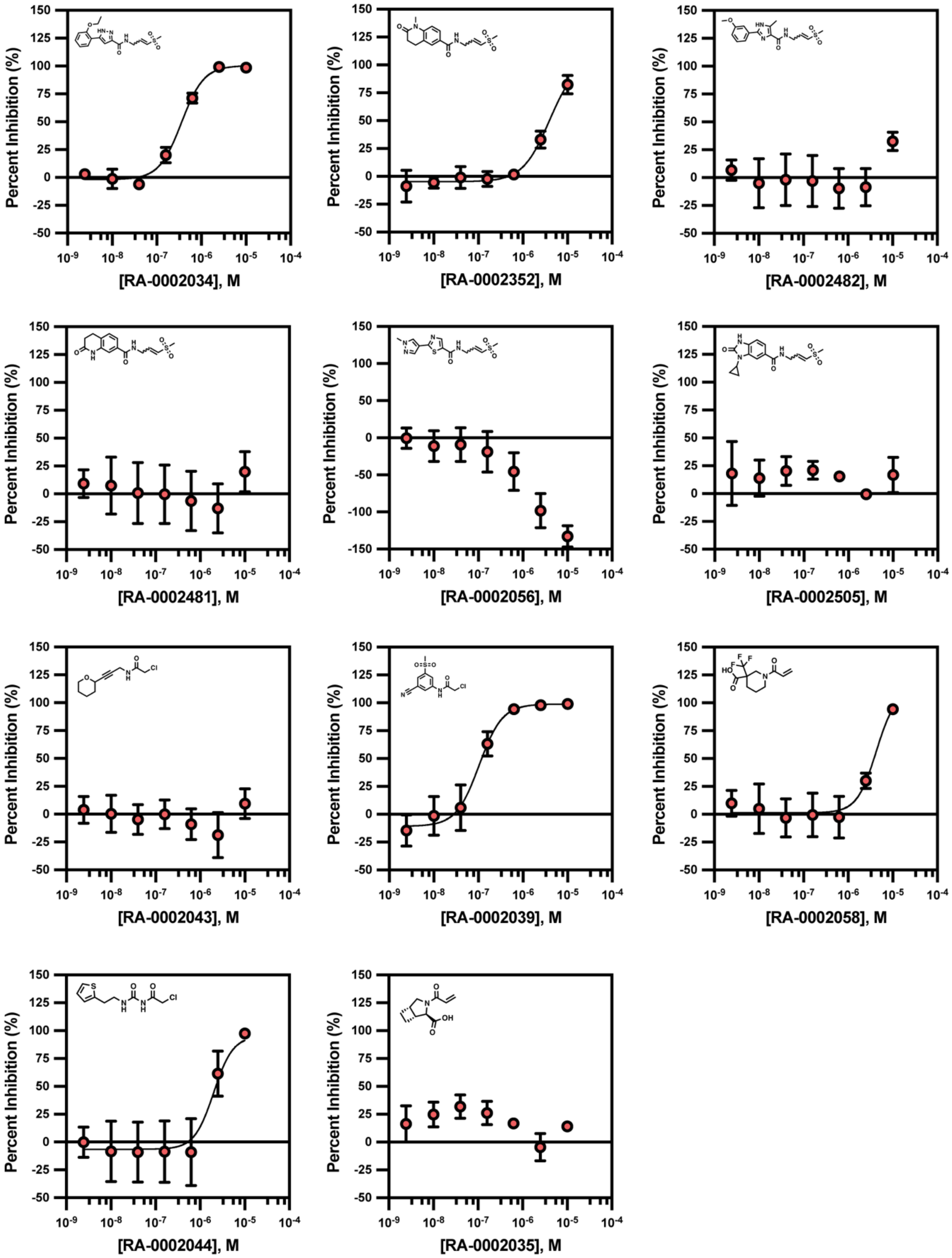


**Figure S17. Covalent fragment VEEV nLuc dose-response results.** Active compounds in CHIKV nsP2pro activity assay were tested in a VEEV nLuc reporter assay in dose-response format, using 4-fold dilutions beginning from 10 μM (n=3).

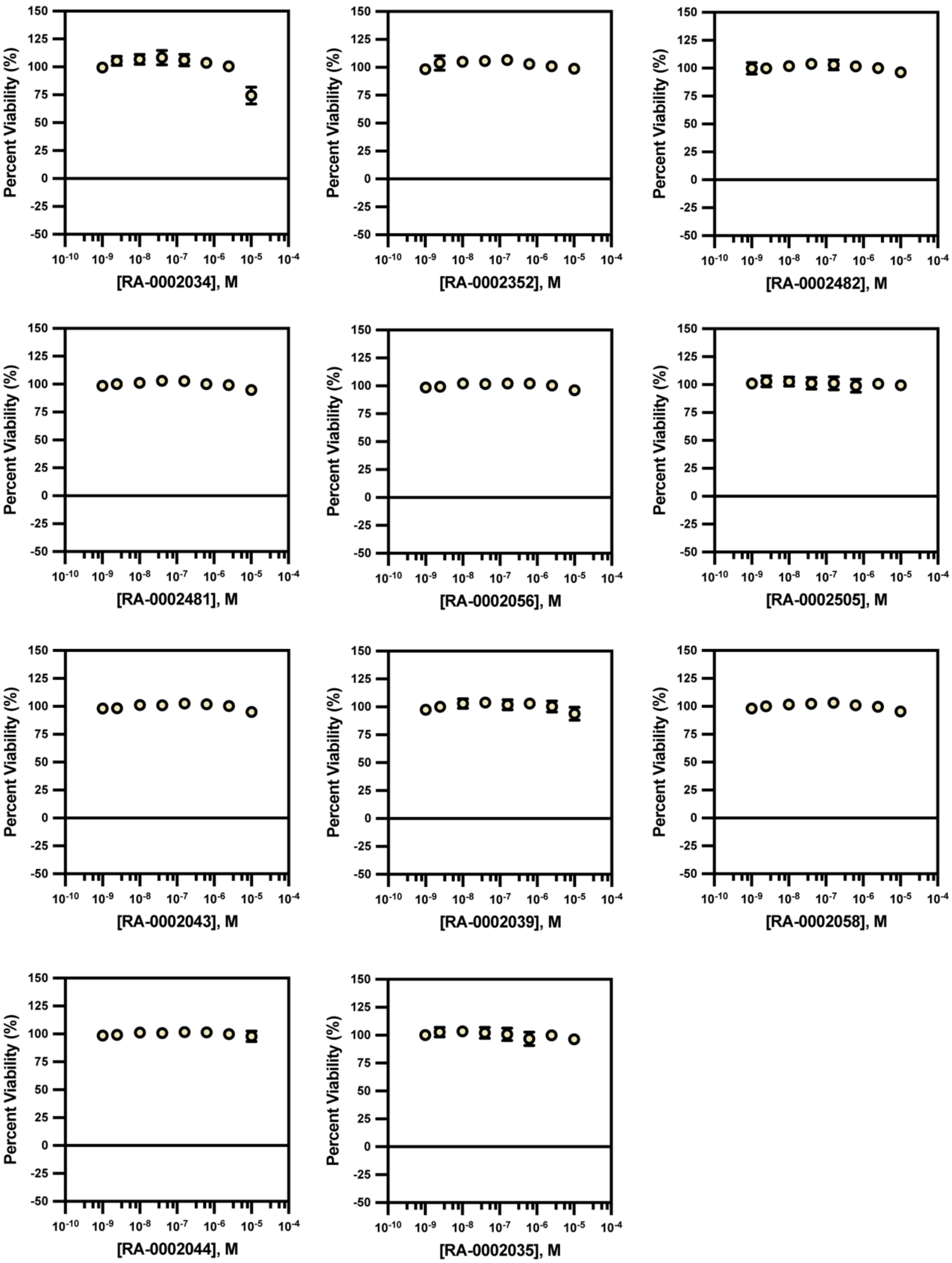


**Figure S18. CTG cell viability.** Active compounds in CHIKV nsP2pro activity assay were tested in a CTG cell viability assay in dose-response format, using 4-fold dilutions beginning from 10 µM (n=3).

**Table S5.** nLuc reporter assay results for covalent fragments.

| Compound | RA ID | Warhead | CHIKV nLuc $IC_{50}$ (μM) | VEEV nLuc $IC_{50}$ (μM) | CTG $IC_{50}$ (μM) |
|---|---|---|---|---|---|
| 85 | RA-0002034 | Internal vinyl sulfone | 0.011 ± 0.004 | 0.32 ± 0.02 | >10 |
| 18 | RA-0002352 | Internal vinyl sulfone | 0.19 ± 0.02 | 3.7 ± 0.8 | >10 |
| 10 | RA-0002482 | Internal vinyl sulfone | 0.61 ± 0.29 | >10 | >10 |
| 16 | RA-0002481 | Internal vinyl sulfone | 3.2 ± 0.3 | >10 | >10 |
| 21 | RA-0002056 | Internal vinyl sulfone | >10 | >10 | >10 |
| 15 | RA-0002505 | Internal vinyl sulfone | 9.8 ± 0.2 | >10 | >10 |
| 11 | RA-0002048 | External vinyl sulfone | | | |
| 46 | RA-0002043 | Chloroacetamide | >10 | >10 | >10 |
| 29 | RA-0002054 | External vinyl sulfone | | | |
| 4 | RA-0002039 | Chloroacetamide | 1.3 ± 0.2 | 0.11 ± 0.02 | >10 |
| 56 | RA-0002057 | Internal vinyl sulfone | | | |
| 30 | RA-0002051 | External vinyl sulfone | | | |
| 70 | RA-0002058 | Acrylamide | 0.61 ± 0.22 | 4.7 ± 1.6 | >10 |
| 66 | RA-0002055 | External vinyl sulfone | | | |
| 51 | RA-0002038 | External vinyl sulfone | | | |
| 107 | RA-0002037 | External vinyl sulfone | | | |
| 124 | RA-0002052 | External vinyl sulfone | | | |
| 99 | RA-0002047 | External vinyl sulfone | | | |
| 83 | RA-0002044 | Chloroacetamide | 4.0 ± 1.2 | 2.1 ± 0.5 | >10 |
| 52 | RA-0002035 | Acrylamide | >10 | >10 | >10 |

**Scheme S4.** Synthesis of RA-0003161 (**154**).

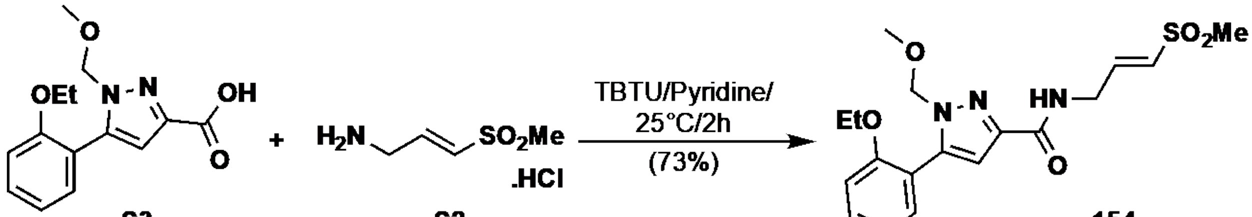


**154**
**RA-0003161**

To a stirred solution of 5-(2-ethoxyphenyl)-1-(methoxymethyl)-1*H*-pyrazole-3-carboxylic acid **S3** (100 mg, 0.36 mmol, 1.0 eq.) and TBTU (174 mg, 0.54 mmol, 1.5 eq.) in pyridine (3.0 mL), (*E*)-3-(methylsulfonyl)prop-2-en-1-amine hydrochloride **S2** (74.5 mg, 0.43 mmol, 1.2 eq.) was added and the reaction was stirred at 25 °C for 2h. On completion of the reaction (based on TLC and LCMS), the reaction was poured into water, washed with aqueous $NaHCO_3$ and brine, and extracted with ethyl acetate. The combined organic layers were dried over anhydrous $Na_2SO_4$, filtered, and concentrated *in vacuo* to give the crude compound. Column chromatography (eluting with 0-100% EtOAc in hexanes), afforded (*E*)-5-(2-ethoxyphenyl)-1-(methoxymethyl)-*N*-(3-(methylsulfonyl)allyl)-1*H*-pyrazole-3-carboxamide **(154)** as a white solid (104 mg, 73%). **$^1$H NMR** (DMSO-$d_6$, 500 MHz): $\delta$ 9.00 (t, *J* = 5.7 Hz, 1H), 7.90 (dd, *J* = 7.7, 1.8 Hz, 1H), 7.46 (s, 1H), 7.33 (ddd, *J* = 8.2, 7.3, 1.8 Hz, 1H), 7.12 (dd, *J* = 8.5, 1.1 Hz, 1H), 7.00 (td, *J* = 7.5, 1.1 Hz, 1H), 6.82 (dt, *J* = 15.3, 4.2 Hz, 1H), 6.74 (dt, *J* = 15.2, 1.6 Hz, 1H), 5.79 (s, 2H), 4.20 – 4.12 (m, 4H), 3.27 (s, 3H), 3.01 (s, 3H), 1.44 (t, *J* = 6.9 Hz, 3H); **$^{13}$C NMR** (DMSO-$d_6$, 126 MHz): $\delta$ 159.2, 155.7, 146.6, 142.9, 136.0, 130.3, 129.5, 127.9, 120.8, 120.5, 112.9, 109.5, 79.9, 63.7, 56.3, 42.2, 38.9, 14.6. (ESI-MS) m/z $[M+H]^+$ 394.

### NMR Spectra

**Figure S19A.** $^{1}$H NMR of RA-0003161 (**154**)

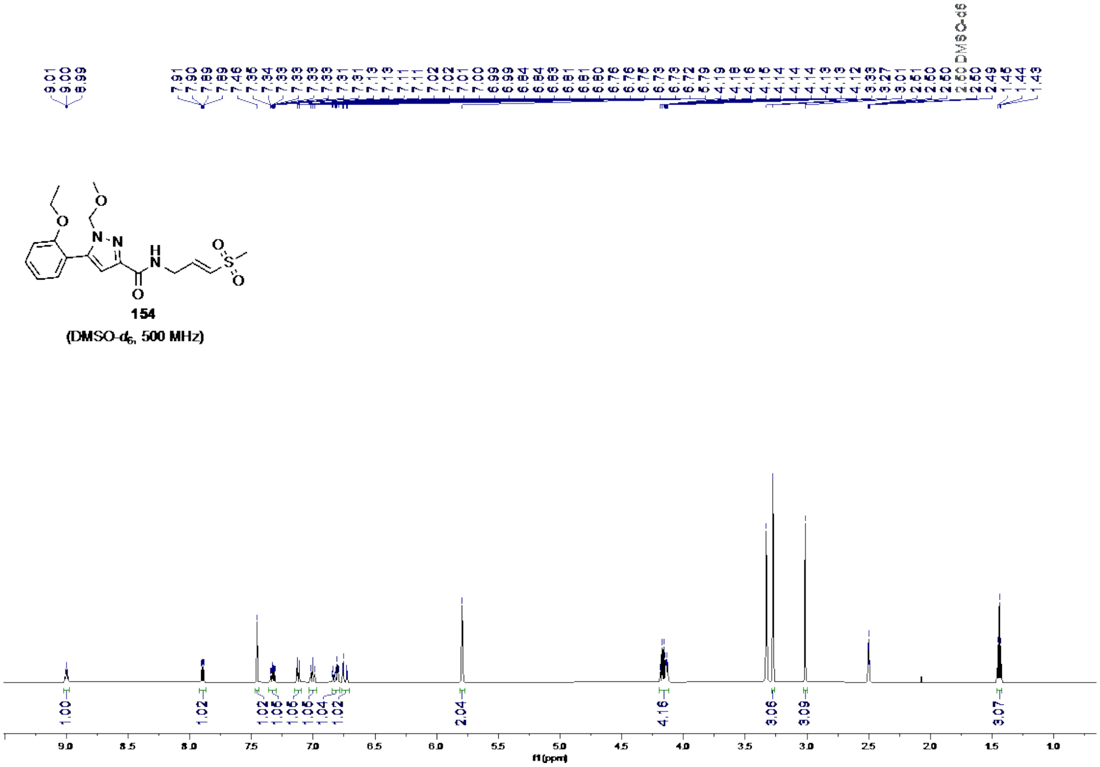


**Figure S19B**. $^{13}$C NMR of RA-0003161 (**154**)

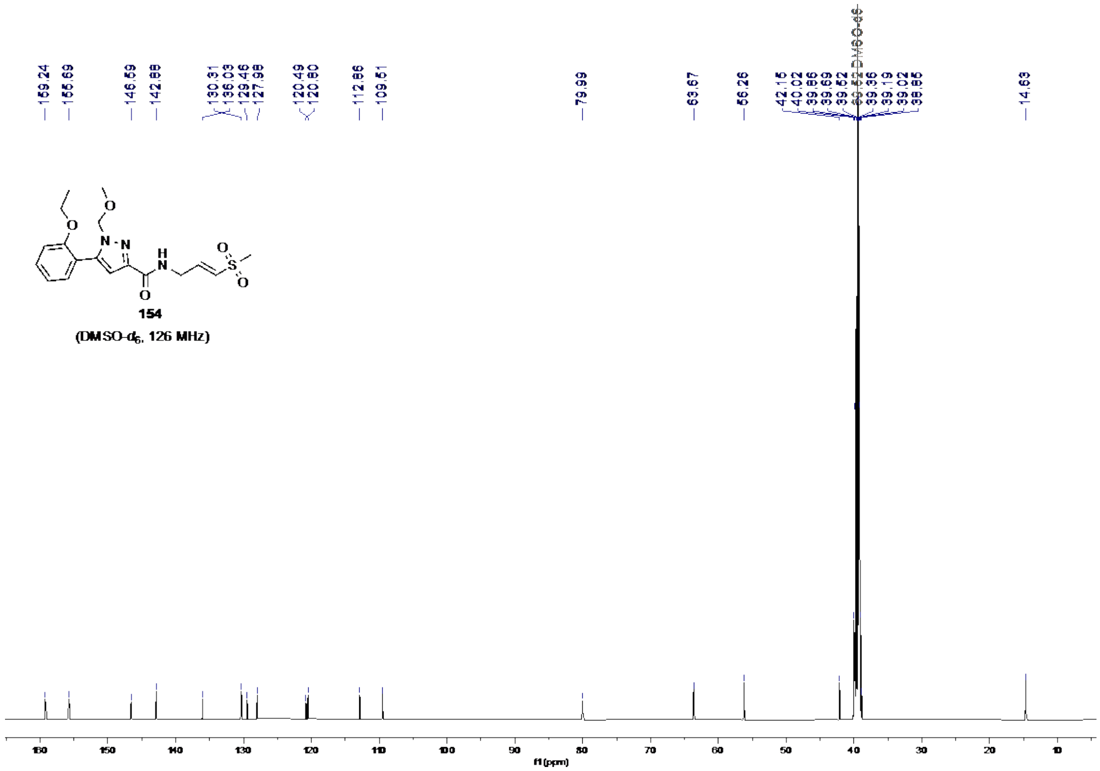

**Scheme S5.** Synthesis of methoxymethyl (MOM)-protected pyrazole carboxylic acid (**S3**)

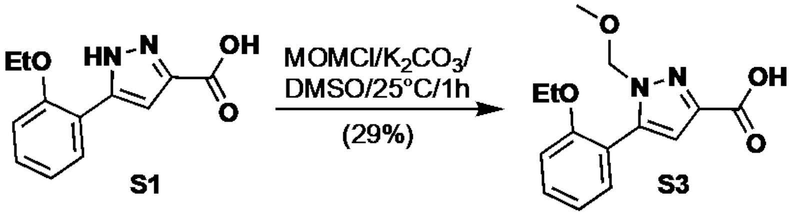


**5-(2-ethoxyphenyl)-1-(methoxymethyl)-1*H*-pyrazole-3-carboxylic acid (S3)**

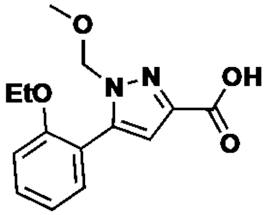


To a stirred solution of 5-(2-ethoxyphenyl)-1*H*-pyrazole-3-carboxylic acid (1.0 g, 4.3 mmol, 1.0 eq.) in DMSO (10 mL), $K_2CO_3$ (1.8 g, 13 mmol, 3.0 eq.) and chloromethyl methyl ether (0.39 mL, 5.2 mmol, 1.2 eq.) were added at 0 °C and the reaction was stirred at 25 °C for 1h. On completion of the reaction (based on TLC and LCMS), the reaction was poured into water and extracted with diethyl ether. The combined organic layers were washed with brine, dried over anhydrous $Na_2SO_4$, filtered, and concentrated *in vacuo* to give the crude compound. Column chromatography (eluting with 10% MeOH in DCM) afforded 5-(2-ethoxyphenyl)-1-(methoxymethyl)-1*H*-pyrazole-3-carboxylic acid **(S3)** as a white solid (350 mg, 29%); $^1$H NMR (DMSO-$d_6$, 500 MHz): $\delta$ 13.52 (s, 1H), 7.93 (dd, *J* = 7.7, 1.7 Hz, 1H), 7.38 (s, 1H), 7.33 (ddd, *J* = 8.9, 7.3, 1.8 Hz, 1H), 7.11 (dd, *J* = 8.4, 1.1 Hz, 1H), 7.01 (td, *J* = 7.5, 1.1 Hz, 1H), 5.77 (s, 2H), 4.15 (q, *J* = 6.9 Hz, 2H), 3.28 (s, 3H), 1.41 (t, *J* = 6.9 Hz, 3H); $^{13}$C NMR (DMSO-$d_6$, 126 MHz): $\delta$ 160.4, 155.8, 146.7, 134.0, 129.6, 127.7, 120.5, 120.3, 112.9, 112.8, 80.2, 63.6, 56.3, 14.7; m/z [M+H]$^+$ 277.

**Figure S20A**. $^{1}$H NMR of compound **S3**

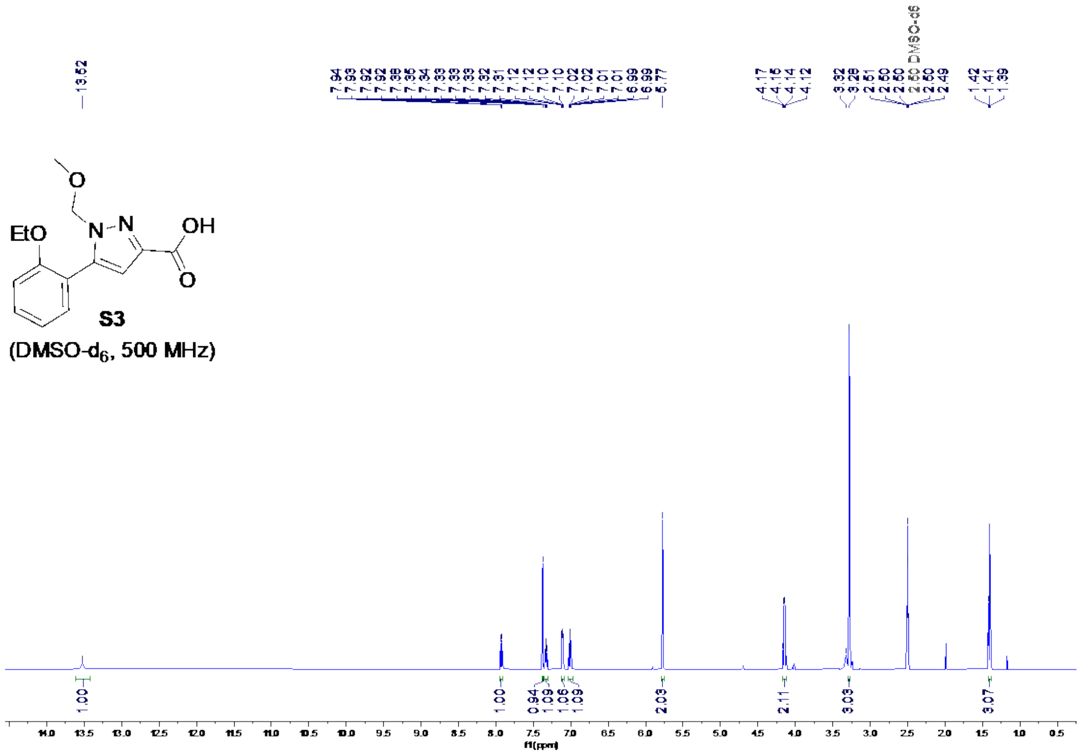

**Figure S20B**. $^{13}$C NMR of compound **S3**

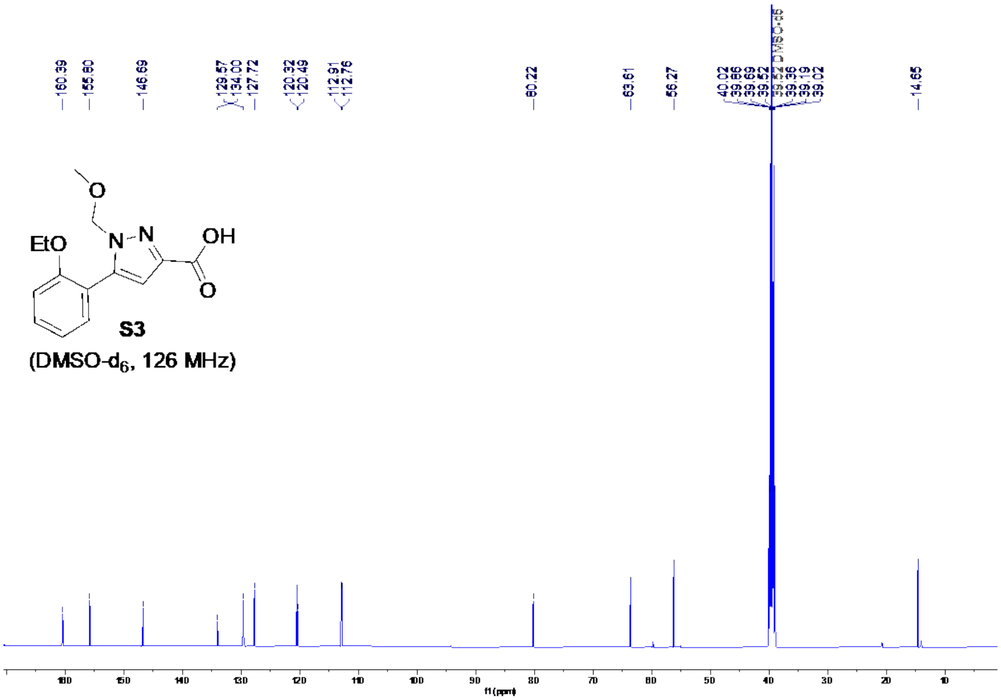

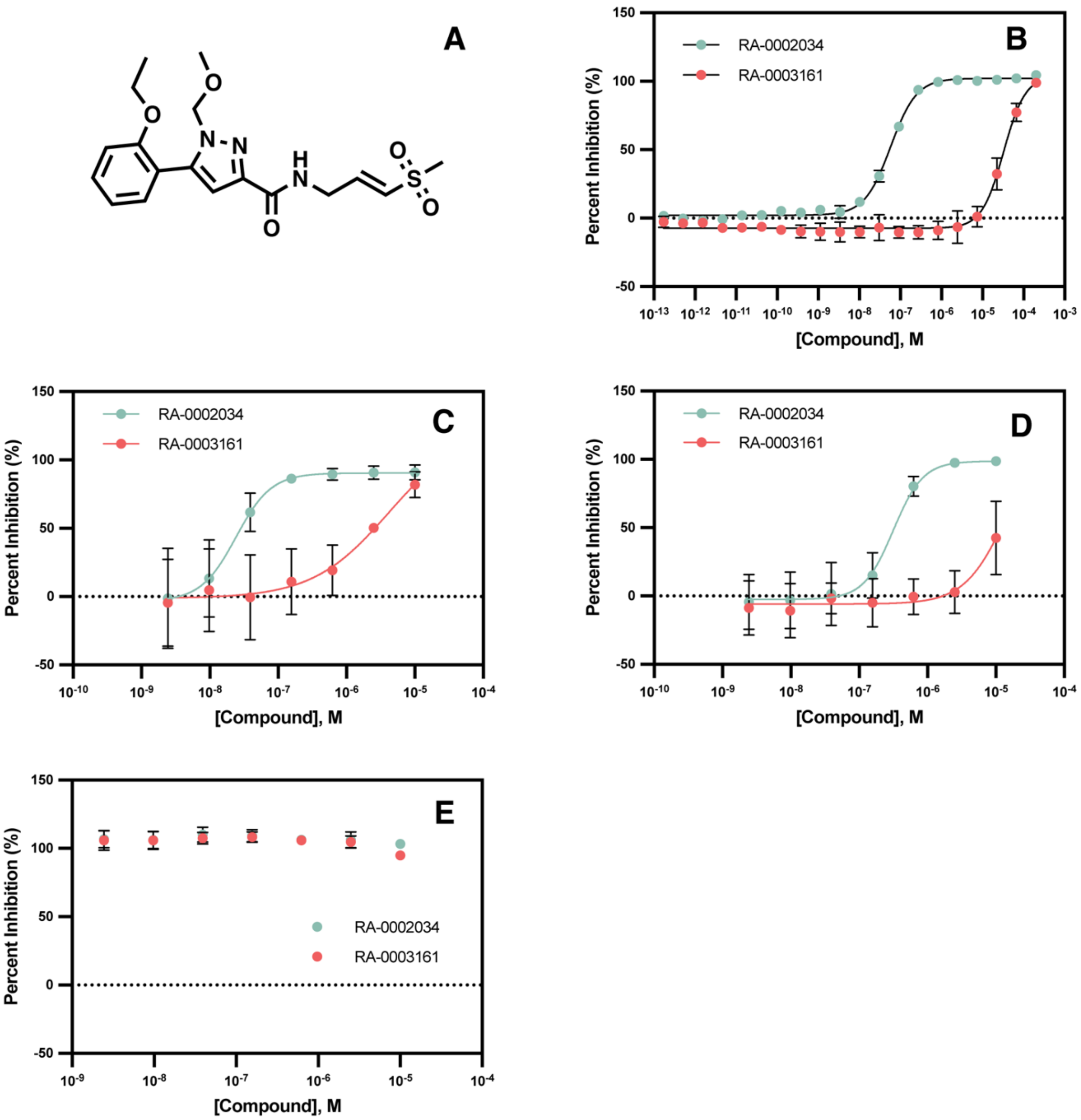


**Figure S21. RA-0003161 inhibition data**. (A) Structure of RA-0003161 (**154**). (B) Dose-response results of RA-0002034 (cyan) and RA-0003161(red) tested in cell-free protease activity assay. Compounds were tested in 3-fold serial dilutions beginning from 200 μM over 20 concentrations. (C) RA-0002034 (cyan) and RA-0003161(red) tested in CHIKV nLuc assay. (D) RA-0002034 (cyan) and RA-0003161(red) tested in VEEV nLuc assay. (E) RA-0002034 (cyan) and RA-0003161(red) tested in CTG assay as a measure of cell viability. For CHIKV nLuc, VEEV nLuc, and CTG assays, compounds were diluted using 4-fold serial dilutions beginning from 10 μM. All curves were fit using the four-parameter Hill equation using GraphPad Prism.

**Supplementary Methods**

**Orthogonal protease activity assays.** Calpain-2 protease was assayed using an internally quenched substrate peptide PLF/AERK conjugated with EDANS and DABCYL on the N- and C- termini respectively. Assay components were diluted in 62 mM imidazole, 0.3 mM $CaCl_2$, 0.10% CHAPS, 0.05% BSA, 1 mM DTT, pH 7.3. Final concentrations were 10 nM calpain-2 and 30 μM of the peptide substrate. RA-0002034 was titrated in a 3-fold serial dilution starting at a final concentration of 200 μM, with a constant DMSO concentration of 2%. TEV protease was assayed using the SensoLyte® 520 TEV Protease Assay Kit according to manufacturer's instructions (Anaspec). Briefly, an internally quenched 7-mer fluorogenic peptide encoding the TEV protease cleavage sequence (ENLYFQG) conjugated with 5-FAM dye and a QXL520 quencher was used as substrate. The 5-FAM fluorophore was excited at 490nm, and emission was read at 520nm. TEV was used at 2 μg/mL and substrate was used at 1X (from 100X stock). RA-0002034 was titrated in a 3-fold serial dilution starting at a final concentration of 500 μM, with a constant DMSO concentration of 5%. Both assays were conducted in a kinetic format and initial velocities in RFU/min were calculated from the slopes of the linear range of the progression curves. Initial velocities were normalized to percent inhibition (%) using fully inhibited enzyme with 10 mM iodoacetamide as 100% inhibition and DMSO only as the uninhibited control. Assays for caspase 1, calpain 1, cathepsin B, cathepsin C, cathepsin H, cathepsin K, cathepsin L, cathepsin S, cathepsin V, papain, SARS-CoV-2 Mpro were conducted in collaboration with Reaction Biology (Table S4).

**MD Simulations.** X-ray structure (PDB ID: 3TRK) were used as the initial structure for the simulations. The MD simulations were performed in the GROMACS 2020.3 package(48, 49), using CHARMM36 force field(50). The protein structure was solvated using the TIP3P water model and neutralized with sodium and chloride ions. Particle mesh Ewald (PME) was used for long-range electrostatic interactions with a 10Å cutoff for non-bonded interactions. The system was initially equilibrated using a NVT thermostat. After that, the system was further equilibrated using NPT thermostats. Next, regular MD simulation was performed at a constant pressure and temperature of 1 atm and 270 K for 500 ns in three replicates. Based on the analysis of the root-mean-square deviation (RMSD) of backbone Ca positions first 50 ns of the simulations were excluded from further analysis to ensure the system equilibration. Real-mean-square-fluctuation (RMSF) values averaged over the three trajectories were used as a measure of conformational flexibility of the protein.

**General Chemistry Methods.** All reactions were performed in oven-dried glassware under an atmosphere of dry N2 unless otherwise stated. All reagents and solvents used were purchased from commercial sources and were used without further purification. $^1$H NMR and $^{13}$C NMR spectra for characterization of new compounds were collected in DMSO-d6 on Bruker 500 MHz and 400 MHz spectrometers at the UNC Eshelman School of Pharmacy NMR Facility. All chemical shifts are reported in the standard notation of parts per million (ppm, δ units) and are referenced to the residual protons in the deuterated solvent used. Coupling constant units are in Hertz (Hz). Splitting patterns are indicated as follows: s (singlet), d (doublet), t (triplet), q (quartet), m (multiplet), dd (doublet of doublets), dt (doublet of triplets), td (triplet of doublets), ddd (doublet of doublets of doublets). Analytical thin layer chromatography (TLC) was performed on pre-coated silica gel plates, 200 μm with an F254 indicator. TLC plates were visualized by fluorescence quenching under UV light or by staining with iodine and KMnO4. Column chromatography was performed using Teledyne ISCO's RediSep Rf® pre-loaded silica gel cartridges on Biotage automated purification systems.

**LCMS Method.** Analytical LCMS data was obtained using a Waters Acquity Ultrahigh-performance liquid chromatography (UPLC) system equipped with a photodiode array (PDA) detector using the following method: solvent A = Water + 0.2% FA, solvent B = ACN + 0.1% FA, flow rate = 1 mL/min. The gradient started at 95% A for 0.05 min. Afterwards, it was ramped up to 100% B over 2 min and held for an additional minute at this concentration, before returning to the initial gradient.

**HPLC Method.** Compounds were purified on preparative HPLC using an Agilent 1100 equipped with a Phenomenex column (PhenylHexyl, 75 x 30 mm, 5 μm) using the following method: Solvent A: water + 0.05 % TFA; Solvent B: MeOH; flow rate: 70.00 mL/min. LC conditions were set at 90 % (A) ramped linearly over 8.0 mins to 100% (B) and held until 10.0 mins at 100% B. At 10.0 mins the gradient was switched back to 90% (A).

**HRMS Method.** HRMS samples were analyzed at the UNC Department of Chemistry Mass Spectrometry Core Laboratory with a Q Exactive HF-X (ThermoFisher, Bremen, Germany) mass spectrometer. Samples were introduced via a heated electrospray source (HESI) at a flow rate of 10 μL/min. HESI source conditions were set as: nebulizer temperature 400 deg C, sheath gas (nitrogen) 20 arb, auxillary gas (nitrogen) 0 arb, sweep gas (nitrogen) 0 arb, capillary temperature 320 degrees C, RF voltage 45 V. The mass range was set to 100-1000 m/z. All measurements were recorded at a resolution setting of 120,000. Solutions were analyzed at 0.1 mg/mL or less based on responsiveness to the ESI mechanism. Xcalibur (ThermoFisher, Breman, Germany) was used to analyze the data. Molecular formula assignments were determined with Molecular Formula Calculator (v 1.3.0). All observed species were singly charged, as verified by unit m/z separation between mass spectral peaks corresponding to the 12C and 13C12Cc-1 isotope for each elemental composition. Separations were conducted on a Waters Acquity UPLC BEH C18 column (2.1 x 50 mm,

1.7 um particle size). LC conditions were set at 95 % water with 0.1% formic acid (A) ramped linearly over 5.0 mins to 100% acetonitrile with 0.1% formic acid (B) and held until 6.0 mins. At 7.0 mins the gradient was switched back to 95% A and allowed to re-equilibrate until 9.0 mins. Injection volume for all samples was 3 uL.
